# Patterns of reduced cortical thickness and striatum pathological morphology in cocaine addiction

**DOI:** 10.1101/306068

**Authors:** Eduardo A Garza-Villarreal, Ruth Alcalá-Lozano, Thania Balducci, Diego Ángeles-Valdéz, M. Mallar Chakravarty, Gabriel A. Devenyi, Jorge J Gonzalez-Olvera

## Abstract

Substance addiction is regarded as an important public health problem, perpetuated by fronto-striatal circuit pathology. A usual finding in neuroimaging human and murine studies is cortical thinning and lower volume when compared to healthy controls. In this study we wished to replicate cortical thinning findings and find if striatum morphology may explain the cortical pathology. For this we analyzed T1w neuroimaging data from an ongoing addiction Mexican dataset. This dataset includes cocaine addicts diagnosed by expert psychiatrists and healthy controls. For the analysis we used voxel-based morphometry, cortical thickness and volumetric analysis of the basal ganglia, and we correlated striatum volume with cortical thickness to find pathological patterns. Our group contrast showed cortical thinning and striatum volume differences in cocaine addicts correlated to their years of substance use, craving and age. Our correlation between striatum-cortex morphology showed higher significant correlations in healthy controls, not observed in cocaine addicts. The correlation between striatum volume and cortical thickness in healthy controls involved similar areas as those shown with less cortical thickness in cocaine addicts. We suggest that striatum morphological changes in addiction may explain the pattern of cortical thinning observed across several substances addiction studies.

**Research Data Related to this Submission:** **Data set** https://zenodo.org/record/1409808#.W5E3oCOZPIF Patterns of reduced cortical thickness and striatum pathological morphology in cocaine addiction

This dataset includes all the data and scripts needed to reproduce the analysis and results on the manuscript “Patterns of reduced cortical thickness and striatum pathological morphology in cocaine addiction” (link). The brain data is not raw, as T1w were not defaced. We will do so in the near future for version 2.0. Instead we include only the “output/thickness” files used in the final analysis. For the use of raw T1w images, please contact the main author EAGV.

## Introduction

Substance addiction is a generalized problem in the world accompanied by important psychiatric comorbidities and exacerbated by poor treatment outcomes ^1^. In Mexico, cocaine is the second most used illegal substance of abuse among the population of users, and it inflicts social, family, economical and health problems (https://www.gob.mx/salud%7Cconadic/acciones-y-programas/encuesta-nacional-de-consumo-de-drogas-alcohol-y-tabaco-encodat-2016-2017-136758). Volkow et al. ^2^ have proposed a hypothesis that substance addiction is a pathological cycle of behavior, and that chronic drug use directly and indirectly affects brain areas such as the striatum and thalamus and their cortical connectivity. Such neurobiological changes perpetuate and potentially reinforce the addiction cycle. The consistent reinforcement of this cycle leads to addictive behaviors commonly observed in individuals suffering from addiction ^3^. As a result, a brain circuit involving the striatum, thalamus and prefrontal areas is commonly studied to better understand how brain function and structure is altered in those suffering from addictions ^4,5^. Recent studies in humans and murine models have supported pathological fronto-striatal connectivity as the main driver of the addiction cycle and substance seeking ^6,7^.

Human neuroimaging studies using voxel-based morphometry (VBM) and cortical thickness (CT) have typically demonstrated variable brain differences, usually including decreased volume or CT in prefrontal, temporal, occipital and subcortical areas in the brains of addicts ^8-13^, although there have been increase volume findings in striatum ^14^. The decrease in these measures is usually correlated with behavioral (i.e. substance craving) and cognitive measures ^15,16^. A meta-analysis in cocaine and methamphetamine addicts found lower volume on bilateral insula, left thalamus, left middle frontal gyrus (lmFG), right anterior cingulate (rACC) and right inferior frontal gyrus (rIFG) ^17^. Parvaz et al ^18^ recently demonstrated that the volume of the inferior frontal gyrus (IFG) and the medial prefrontal cortex (mPFC) increase following treatment for cocaine addiction. These studies suggest brain morphology is highly affected in addiction, and that treatment success can partly revert this damage. These areas found with reduced volume or CT in addiction such as the insula, IFG, mPFC, ACC, have also been found connected structurally and functionally to the different nuclei of the striatum ^19^. If striatum is a region highly affected in addiction and it is intimately connected with these cortical areas, which in turn seem to be commonly found affected in addiction, pathological morphological changes in striatum may help explain the reduction of cortical thickness and volume in such widespread areas of the cortex.

In this study, we wish to find neuroanatomical differences between cocaine addicts and matched healthy controls, to better understand the toxic effect of drug use and the effect of craving. We also wanted to study the relation between striatum structure and cortical thickness. For this, we used novel computational anatomy algorithms to perform volumetric analysis, cortical thickness extraction and subcortical segmentation of striatum and thalamus.

## Materials and Methods

### Participants

We recruited 160 participants as part of a principal multidisciplinary addiction study at the Instituto Nacional de Psiquiatría “Ramón de la Fuente Muñiz” in Mexico City, Mexico. Of those, 49 participants did not fulfill the inclusion criteria. We diagnosed cocaine dependence using the MINI International Neuropsychiatric Interview Spanish version ^20^, which was administered by trained psychiatrists. For inclusion, cocaine consumption had to be active or with abstinence less than 60 days prior to the scan, with frequency of use of at least three days per week and no more than 60 continued days of abstinence during the last 12 months. There could be polysubstance use, however cocaine had to be the drug of impact. Additional exclusion criteria for both groups were: somatic diseases, neurological disorders, severe suicidal risk, history of head trauma with loss of consciousness, pregnancy, obesity, severe psychiatric disorders and non-compliance with magnetic resonance imaging safety standards. A final sample of 64 cocaine addicts (AD) (7 female) and 47 healthy controls (HC) (8 female) were included in our study. Healthy controls were matched as closely as possible by age (± 2y), sex and handedness. Education was matched as closely as possible, though it has significantly higher in HC, therefore education was added as a covariate in the statistical analysis. Table 1 describe the demographic and addiction related information. The study was approved by the local ethics committee and performed at the Instituto Nacional de Psiquiatría “Ramón de la Fuente Muñiz” in Mexico City, Mexico. The study was carried out according to the Declaration of Helsinki. All participants were invited through posters placed in several centers for addiction treatment and through the Institute’s addiction clinic for outpatients. Healthy controls were recruited from the Institute (i.e. administrative workers, their family, etc) and using Internet social outlets. Participants provided verbal and written informed consent. The participants underwent clinical and cognitive tests besides the MRI as part of the main ongoing addiction database. Participants were asked to abstain from drug use for at least 24 hours prior to the study and were urine-tested for the presence of the drugs and a breath determination of alcohol in the blood before the MRI scan. The clinical, cognitive and MRI sessions were performed either the same day as minimum, or 4 days apart as maximum. It is important to point out none of our participants were homeless or in extreme poverty.

**Table 1.** Demographic and substance addiction variables between groups.

| Variable | HC<br>(N=47) | AD<br>(N=64) | p |
| --- | --- | --- | --- |
| Age | 30.7 ± 7.6 | 31.0 ± 7.2 | 0.846 |
| Sex |  |  | 0.519 |
| - F | 8 (17.0%) | 7 (10.9%) |  |
| - M | 39 (83.0%) | 57 (89.1%) |  |
| Education | 3.7 ±1.5 | 2.9 ±1.2 | 0.003 |
| Handedness |  |  | 0.99 |
| - A | 3 ( 6.4%) | 4 ( 6.2%) |  |
| - L | 4 ( 8.5%) | 5 ( 7.8%) |  |
| - R | 40 (85.1%) | 55 (85.9%) |  |
| BIS Total | 40.6 ±11.5 | 60.9 ±15.2 | < .001 |
| • BIS ICog | 11.9 ±3.7 | 17.2 ± 5.4 | < .001 |
| • BIS IMo | 12.7 ±6.0 | 18.2 ±7.5 | < .001 |
| • BIS INoPI | 15.9 ±5.6 | 25.5 ±7.6 | < .001 |
| Cigarettes per day (tobacco) | 1.2 ±1.4 | 3.7 ±4.4 | 0.006 |
| Initial Age | - | 21.67 ±6.15 | - |
| Years of Consumption | - | 9.3 ±6.58 | - |
| Craving (CCQ General) | - | 140 ±42.49 | - |

HC = Healthy Control; AD = Cocaine Addict; F = female; M = male; A = Ambidexterous; L = Left handed, R = Right handed, BIS = Barrat Impulsivity Scale; ICog = Cognitive; IMo = Motor; INoPI = Non-Planning; CCQ = Cocaine Craving Quotient.

### Clinical measures

Craving in the last month and at the interview was measured using the cocaine craving questionnaire (CCQ) ^21^. Self-reported impulsivity was evaluated with the Barratt Impulsiveness Scale (BIS-11), which has three subscales: non-planning impulsiveness, which involves a lack of forethought; cognitive impulsivity, which involves making quick decisions; and motor impulsivity, which involves acting without thinking ^22^.

### MRI Acquisition

T1-weighted brain data were acquired using a Philips Ingenia 3T Magnetic Resonance Imaging (MRI) system (Philips Healthcare, Best, Netherlands & Boston, MA, USA) with a 32-channel dS Head coil. T1-weighted images were acquired using a 3D FFE SENSE sequence, TR/TE = 7/3.5 ms, FOV = 240, matrix = 240 x 240 mm, 180 slices, gap = 0, plane = Sagittal, voxel = 1 x 1 x 1 mm (5 participants were acquired with a voxel size = .75 x .75 x 1 mm), scan time = 3.19 min. As part of the principal addiction database, resting state fMRI, High Angular Resolution Diffusion Imaging (HARD), and Diffusion Kurtosis Imaging (DKI) sequences were also acquired and are not part of this study. The order of the sequences was: rsfMRI, T1w, HARDI, DKI, and was maintained across participants. Total scan time was ∼50 minutes.

### Image preprocessing and processing

T1-weighted images were converted from DICOM format to MINC for preprocessing. T1 images were preprocessed using an in-house preprocessing pipeline with the software Bpipe (https://github.com/CobraLab/minc-bpipe-library)^23^, which makes use of the MINC Tool-Kit (http://www.bic.mni.mcgill.ca/ServicesSoftware/ServicesSoftwareMincToolKit) and ANTs ^24^. Briefly, we performed N4 bias field correction ^25^, linear registration to MNI-space using ANTs, we cropped the region around the neck in order improve registration quality, followed by transformation back to native space, and created whole-brain masks.

We estimated volume-based (VBM) and surfaced-based variables (cortical thickness [CT] and surface area [SA]) using the CIVET processing pipeline (version 1.1.12; Montreal Neurological Institute). First, the T1w images were linearly aligned to the ICBM 152 average template using a 9-parameter transformation (3 translations, rotations, and scales) ^26^ and preprocessed to minimize the effects of intensity non-uniformity ^27^. The images were then classified into three main tissues: gray matter (GM), white matter (WM) and cerebrospinal fluid (CSF) ^28^. GM was used for VBM. The hemispheres were modeled as GM and WM surfaces using a deformable model strategy that generates 4 separate surfaces defined by 40962 vertices each ^29^. CT was derived between homologous vertices on GM and WM derived using the t-link metric and blurred with a 20 mm surface-based diffusion kernel, while SA was estimated by averaging across the adjoining faces at each vertex ^30^. Native-space thicknesses were used in all analyses reported ^31,32^. Homology across the population was achieved using a non-linear surface-based normalization that utilizes a mid-surface (between pial and WM surfaces) ^33^. This normalization uses a depth-potential function ^34^ that fits each subject to a minimally biased surface-based template ^35^.

For the subcortical analysis, the native space preprocessed files were input into the MAGeT-Brain morphological analysis pipeline (http://cobralab.ca/software/MAGeTbrain.html) ^36^. MAGeT-Brain is modified multi-atlas segmentation technique designed to take advantage of hard-to-define atlases and uses a minimal number of atlases for input into the segmentation process. The used a basal ganglia atlas ^37^ obtained by manual segmentation of one brain. We obtained segmentation and volume measures for striatum, thalamus and globus pallidus.

### Statistical analysis

Voxel based morphometry (VBM) gray matter and vertex-wise analyses were performed with the RMINC package (https://wiki.phenogenomics.ca/display/MICePub/RMINC) in R statistics and RStudio ^38^. Public packages used for the analysis were: tidyverse, psych, pastecs, moonBook and plotrix. The general linear model included “CT” as the dependent variable, “group” as the between subjects variable, and “age”, “sex” and “education” as covariates. All analyses were corrected for multiple comparisons using the false discovery rate (FDR) at 10% ^39^. From the resulting significant peaks we extracted MNI coordinates and labels based on the AAL atlas ^40^, except for VBM in which we used Harvard-Oxford Cortical Atlas ^41-44^. As post-hoc, we calculated the correlation coefficient between years of consumption and craving, and all the significant peaks CT. Using that matrix, we statistically analyzed only correlations that exceeded a chosen threshold of r = ± 0.2 (low-medium effect size) using the t-distribution with an alpha of 0.05. We then used the FDR to adjust the p-value for multiple comparisons of the correlations. As a side note, VBM was only calculated because it is a more widely used measure of brain morphology. Because we were more interested in cortical thickness, we did not further analyze this measure. However, tools such as BrainMap (http://brainmap.org) would be able to use this data for future meta-analyses.

### Basal ganglia analysis

We studied basal ganglia volumes using a general linear model that included subcortical volume as the dependent variable, group as the between-subjects variable, and age, sex and education as covariates. Our previous study in cocaine addiction showed mainly group x age interactions in striatum volume ^45^, hence we performed that interaction model as well. We then calculated the correlation coefficient between years of consumption and craving, and all basal ganglia including their striatum and thalamus segmentation. Using that matrix, we statistically analyzed only correlations that exceeded a chosen threshold of r = ± 0.2 (low-medium effect size) using the t-distribution with an alpha of 0.05. Because basal ganglia volume is relative to whole-brain volume, we then performed partial correlations controlling for whole-brain volume.

### Striatum-cortex correlation analysis

The covariation between striatum subnuclei that were correlated significantly with years of consumption and craving (left nucleus accumbens and right precommissural precuneus), and whole-brain cortical thickness was studied using a similar approach to the “Mapping anatomical correlations across cerebral cortex (MACACC)” analysis method ^46^. The MACACC method is performed by selecting a seed region of interest (ROI) and correlating the CT of this ROI with the CT of all brain vertices. This approach is similar to functional connectivity analysis ^47,48^. The resulting statistic gives an indication of the degree to which CT throughout the brain covaries with the ROI across subjects and can be used to estimated the structural and functional connectivity between different areas. As our ROI, we chose to instead use the volume of the significant nuclei in the partial correlation analysis: left nucleus accumbens and right pre-commissural putamen volume (basal ganglia analysis). We also chose whole left and right striatum volume to corroborate our results. This method has been used successfully ^49^. We then correlated the volume of each ROI against the brain vertices for all participants, and then each group separately. All maps were FDR corrected at 5% due to the high distribution of significant peaks.

## Results

We found significantly lower volume (Supplementary Figure 1 & Table 1) and cortical thickness in cocaine addicts in mainly prefrontal areas (Figure 1 and Supplementary Table 2). VBM showed 2 small clusters of increased volume that were not found in the CT analysis. Surface area was not significant.

**Figure 1.**
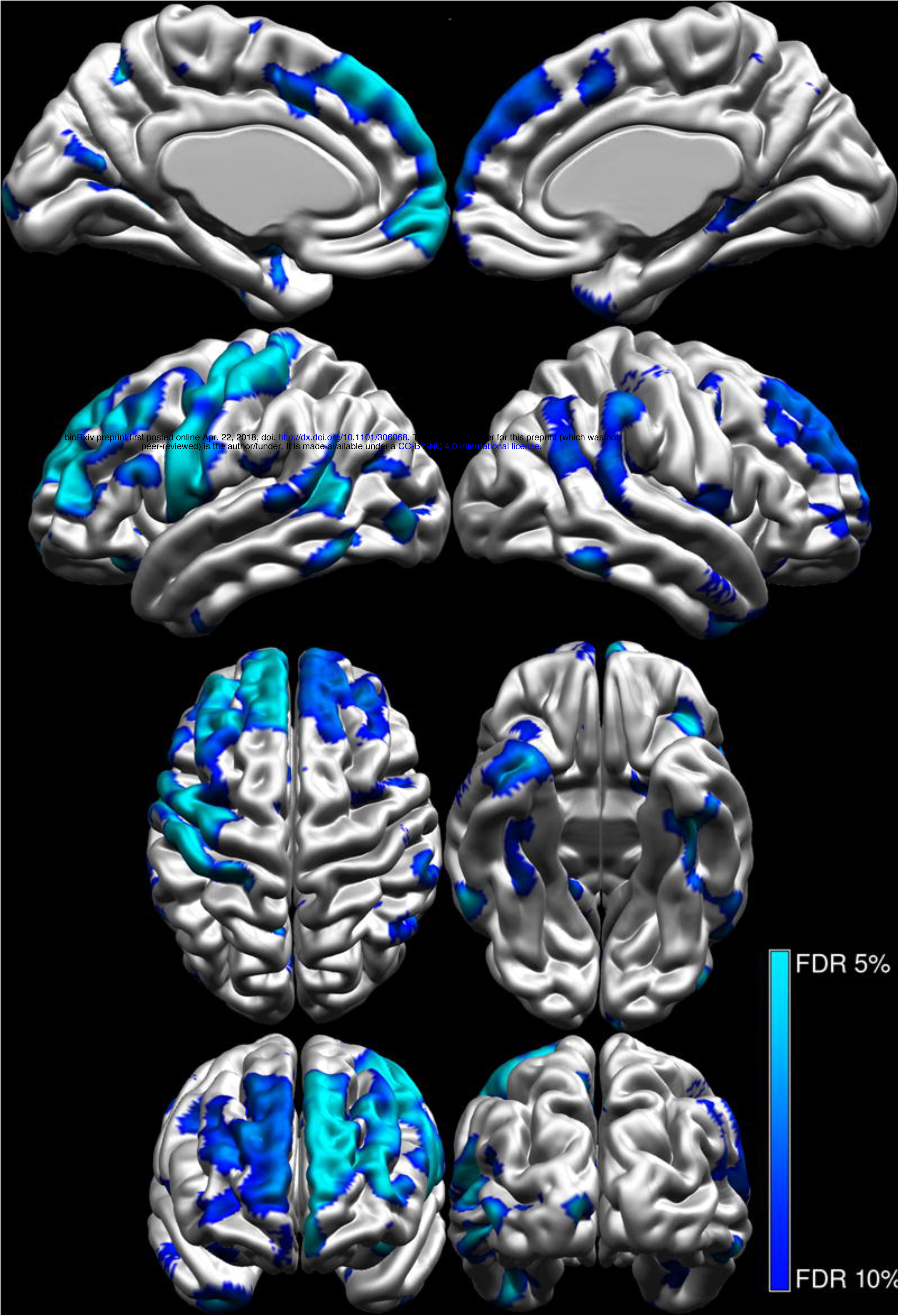
Cortical thickness group difference. Left column views = 1) medial, left hemisphere, 2) lateral, left hemisphere, 3) superior, 4) frontal. Right column views = 1) media, right hemisphere, 2) lateral, right hemisphere, 3) inferior, 4) occipital.

The post-hoc analysis showed that years of cocaine consumption and craving were significantly correlated with several CT peaks (Supplementary Figs. 2 and 3, Supplementary Tables 2 and 3). The subcortical analysis showed a significant lower volume in left thalamus of the AD group (F (1,104) = 4.723, p = 0.03) (Supplementary Fig. 4) than the HC group. There was no difference in striatum and globus pallidus volume. There was, however, a significant group x age interaction in left and right striatum volume (Figure 2) at alpha 0.1 (left: F (1,103) = 2.83, p = 0.1; right: F (1,103) = 3.26, p = 0.07), which was similar to our previous findings ^45^.

**Figure 2.**
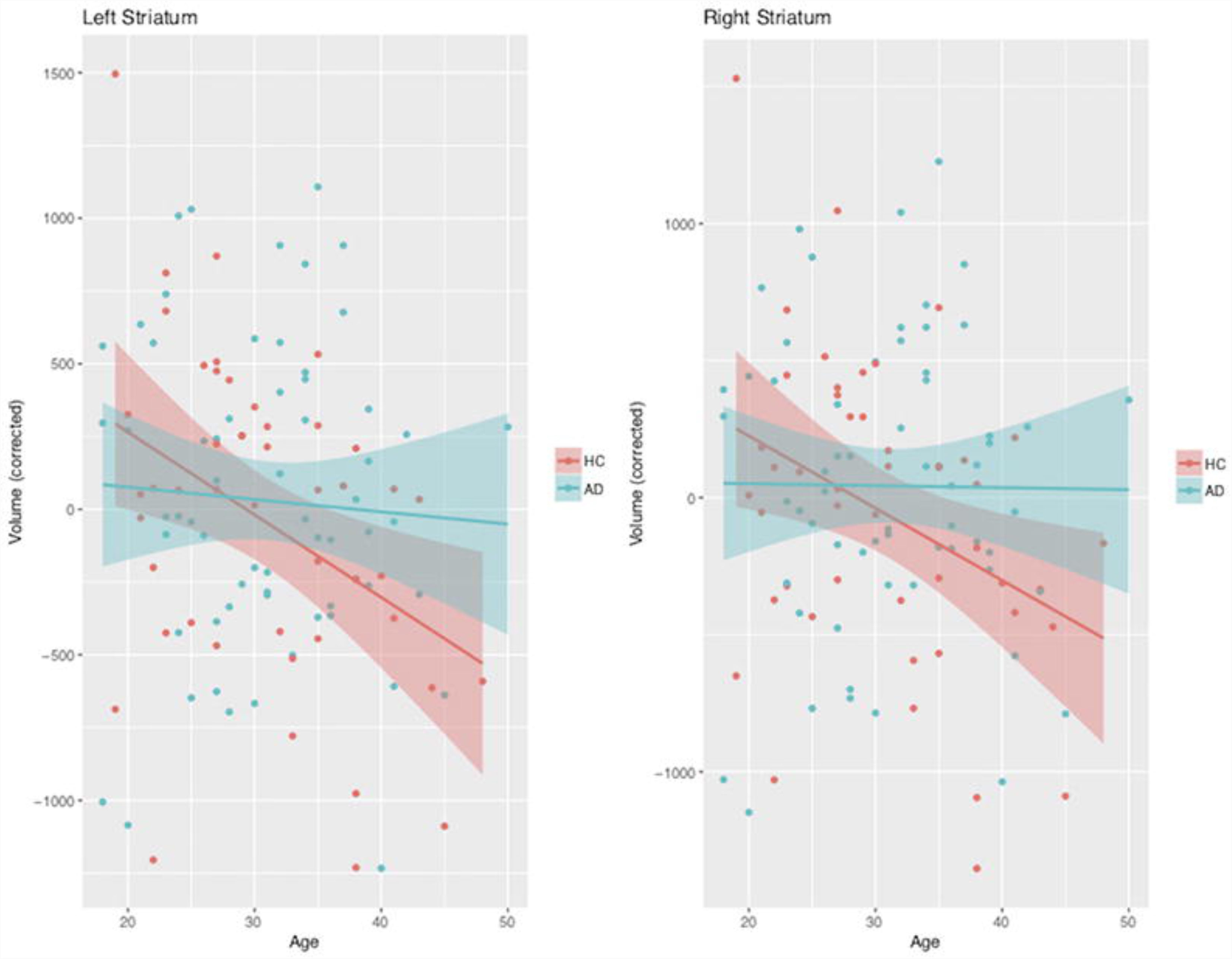
Scatter plot of group x age interaction in striatum volume. HC = Healthy controls; AD = cocaine addicts. The corrected volumes are the residuals of the linear model without group and age.

The bivariate correlation analysis of years of consumption and craving with basal ganglia volumes, showed no significant results. The partial correlation analysis controlling for whole-brain volume showed significant negative correlation between: 1) years of consumption and left nucleus accumbens (r = −0.23, t = −2.46, p = 0.02), and 2) craving and right pre-commissural putamen (r = −0.27, t = −2.89, p = 0.005).

The striatum-cortex correlation analysis results using the ROIs: 1) left nucleus accumbens (lNAcc) and 2) right pre-commisual putamen (rPrePut), are shown in Figure 3. The result of left and right whole striatum volume are in the Supplementary Figure 5. The resulting significant covariance maps show lNAcc and rPrePut volumes are related to similar areas that showed lower cortical thickness in AD in the CT group comparison (Figure 1). A subset analysis showed that the correlation between lNAcc and cortex in AD is nonexistent compared to HC. The subset analysis of correlation between rPrePut and cortex showed higher significant correlations in HC than the AD group. For this last analysis, significant brain areas of correlation shared between groups are shown in Table 2. As for whole striatum volume, we also found higher correlation in HC compared to AD. All peak tables are shown in Supplementary Tables 5 to 15. In general, striatum volume in AD showed low correlation to cortical thickness compared to healthy controls.

**Figure 3.**
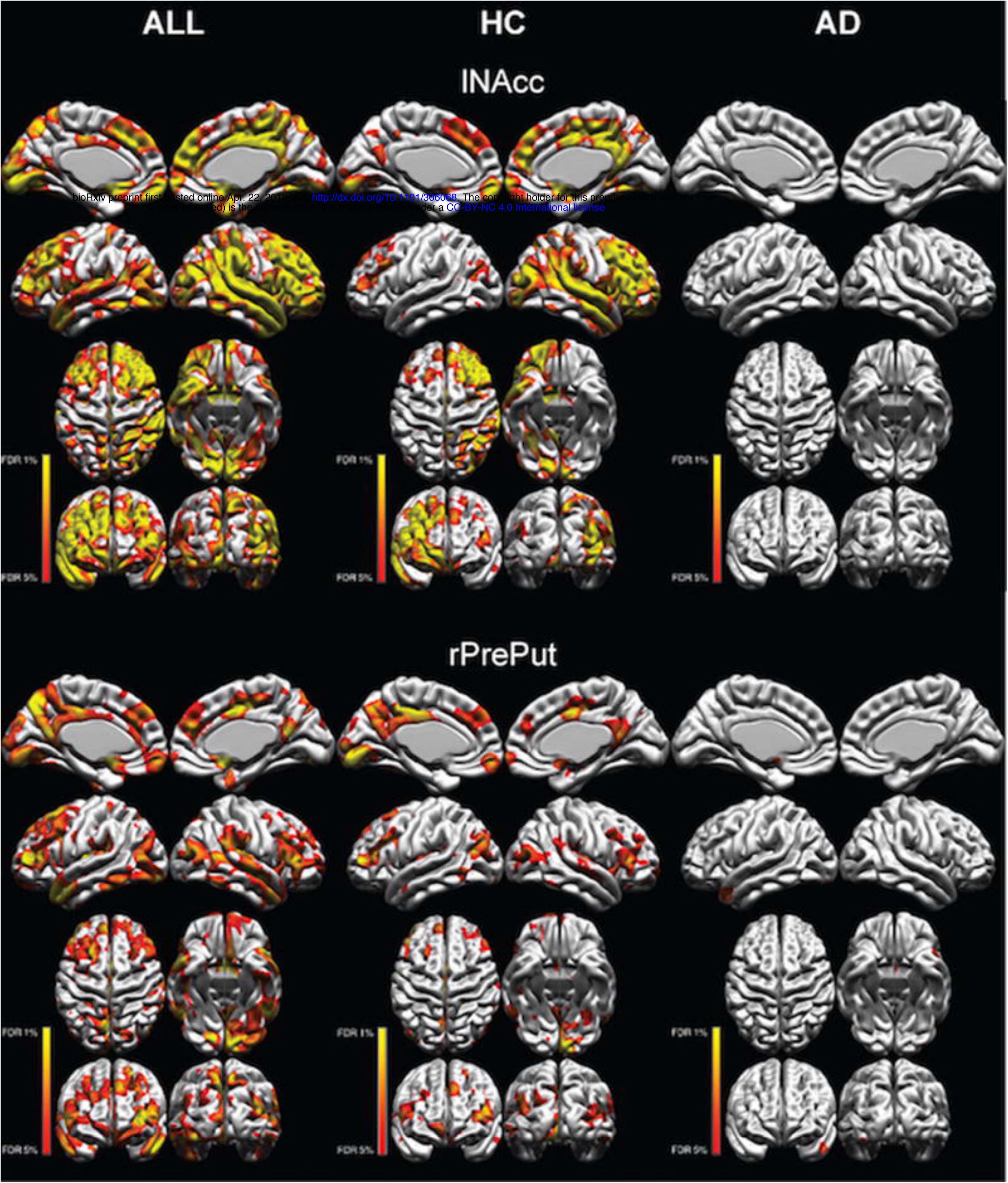
Correlation of left nucleus accumbens and right pre-commissural putamen with cortical thickness. Red-yellow colours show significant vertices. HC = Healthy controls, AD = cocaine addicts, lNAcc = left nucleus accumbens, rPrePut = right pre-commissural putamen. In lNAcc, for the AD group, there were were no significant peaks below FDR 5%.

**Table 2.** Significant peaks in similar brain areas between groups in the correlation analysis between right precommissural putamen and cortical thickness.

| Brain Area | HC |  |  |  |  | AD |  |  |  |  |
| --- | --- | --- | --- | --- | --- | --- | --- | --- | --- | --- |
|  | vertex | x | y | z | t | vertex | x | y | z | t |
| Left Superior temporal gyrus | 10435 | -41 | -18 | -1 | 3.28 | 13033 | -44 | 3 | -15 | 4.86 |
| Left Middle temporal gyrus | 35910 | -61 | -54 | -7 | 3.75 | 13282 | -49 | 13 | -29 | 4.36 |
| Left Inferior occipital gyrus | 576 | -59 | -60 | -9 | 3.92 | 33470 | -36 | -88 | -17 | 4.18 |
| Left Precentral gyrus | 26769 | -24 | -3 | 54 | 2.88 | 20887 | -53 | -5 | 30 | 3.94 |

HC = healthy controls, AD = cocaine addicts, vertex = Surface vertex CIVET 1.1.12, t = tvalue.

## Discussion

In our study, Mexican cocaine addicts showed lower gray matter volume and cortical thickness in several brain areas, with the most extensive difference on prefrontal cortex. Cortical thickness and striatal subnuclei volume were significantly correlated to years of cocaine use and craving. The covariation between striatal whole and subnuclei volumes, and cortical thickness suggests close neuroanatomical pathology of fronto-striatal areas in cocaine addicts.

Using VBM and CT analysis we found significantly lower values in cocaine addicts (AD) than healthy controls (HC). In VBM we found increased volume in two areas that were not found in the CT analysis, probably due to the differences between methods. These findings of cortical thinning are not surprising as they have been shown in other studies of different type of addiction ^9,10,50^, corroborating cortical pathology from chronic cocaine use. The cortical thinning was observed in areas of all cortical lobes, slightly lateralized to the left hemisphere. Although we did not find group differences between volumes in striatum, we found an age x group interaction that suggests a pathological development related to addiction and or chronic use. The striatal subnuclei nucleus accumbens and precommissural putamen were correlated with years of cocaine use and craving. Striatum volume differences, when compared to healthy controls, can be present or absent, higher or lower, in different studies ^14,51^, which suggests either a complex pathology when measured with these methods or differences in volumetric and segmentation methods. Nevertheless, animal and humans studies corroborate the involvement of the striatum in addiction and affectation of its morphology ^52-54^.

A landmark study by Chen et al ^6^ using optogenetics showed that seeking for cocaine in addicted mice could be effectively stopped by stimulating the prelimbic cortex (mPFC or DLPFC in human), while inhibiting this region induced the opposite effect, increased cocaine seeking. In humans, a similar effect has been observed by stimulating the DLPFC using rTMS where cocaine addicts report reduced craving and cocaine use ^55^. The fronto-striatal circuit is involved in response inhibition, which is found to be greatly affected in cocaine addiction in animal models and in human studies, and this circuit includes the striatum, thalamus, globus pallidus, primary motor cortex, ACC, dmPFC and the vlPFC ^5^. However, the structural connectivity between the striatum and cerebral cortex seems to be more extended, involving also areas such as the SFG, IFG, temporal pole and occipital cortex ^19^. Substance addiction is a complex condition and at the moment the main hypothesis for the etiology of the addictive cycle is the fronto-striatal circuit pathology ^2,56^. Although the causal direction of the pathology (fronto-striatal or striato-frontal) has not been demonstrated yet, the involvement of dopamine receptors in striatum suggests a mainly striatal pathology ^57^.

The covariation between lNAcc/rPrePut volume and CT of all participants (striatum-cortex covariation) seems to follow a similar pattern to the structural connectivity of the striatum. Interestingly, not only our group contrast map of lower CT in cocaine addicts shows similarities with the striatum-cortex covariation, but also this covariation was only observed in healthy controls and it almost disappeared in cocaine addicts. This could be an indication of the underlying pathological changes in striatum or cortex (or both), secondary to cocaine addiction or chronic use. This shared morphological finding have been shown in young adult smokers with lower CT in frontal cortical areas and higher volume of the caudate ^58^. In a study with several types of substances, another study showed lower volume in frontal areas as well as the caudate nucleus, among others ^59^. As for the involvement of striatum pathology and cortex morphology in human addiction, a recent study showed that striatal D1-type receptor (dopamine) levels are correlated with mean global cortical thickness in methamphetamine users but not in controls, specifically in temporal and occipital lobes ^60^. The authors suggest this abnormality may be a cortical adaptation to chronic substance use with involvement of the D1-type. The evidence highly suggests that the observed morphological findings in cortex may be due to either striatum pathology, or cortical pathology may drive the striatum changes that engrain this pathology. Confirming a causal relationship would help explain the shared cortical findings across types of substance addiction and would corroborate the hypothesis about dopamine related fronto-striatal dysfunction as one of the main causes of human substance addiction.

Our study has several limitations. Correlational studies cannot prove causality as they can only suggest relationships that can be studied further in real experimental designs. However, it is obvious that experimental designs in substance addiction are unethical in humans; hence we rely on animal studies and correlational designs in humans to provide knowledge. Our significant threshold for the multiple comparisons FDR in the CT analysis was 10% (q = 0.1), which may be considered more liberal than usual and caution should be taken when interpreting our findings. However, this approach was preferred to allow for a more exploratory study and we have successfully used it in our previous studies. Our dataset is unique and ongoing, and because addiction and polysubstance use is complex, we wanted to avoid false negatives. The corroboration of our results in relation to other studies seem to support our use of a more relaxed threshold in this particular sample. Our sample is mainly males due to the prevalence of cocaine addiction in this sex, which is a problem in all cocaine addiction studies.

Our results show a possible relation between striatum volume and cortical thinning in cocaine addiction, which further confirms fronto-striatal pathology. Specifically, we believe our results suggest that the pattern of cortical thinning found in most addiction studies may be explained by striatal pathology. Future studies should aim at corroborating the cortical connectivity between striatum and cortex in substance addiction using advance non-invasive diffusion methods.

## Acknowledgements

We would like to thank Rocio Estrada Ordoñez and Isabel Lizarindari Espinosa Luna at the Unidad de Atención Toxicologica Xochimilco for all their help and effort. Finally, we thank the study participants for their cooperation and patients. This project was funded by CONACYT-FOSISS-S0008 project No.0260971, No.0201493 and CONACTT-Catedras project No.2358948. For the use of the Harvard-Oxford Atlas, we are very grateful for the training data for FIRST, particularly to David Kennedy at the CMA, and also to: Christian Haselgrove, Centre for Morphometric Analysis, Harvard; Bruce Fischl, Martinos Center for Biomedical Imaging, MGH; Janis Breeze and Jean Frazier, Child and Adolescent Neuropsychiatric Research Program, Cambridge Health Alliance; Larry Seidman and Jill Goldstein, Department of Psychiatry of Harvard Medical School; Barry Kosofsky, Weill Cornell Medical Center.

The study was approved by the local ethics committee and performed at the Instituto Nacional de Psiquiatría “Ramón de la Fuente Muñiz” in Mexico City, Mexico. The study was carried out according to the Declaration of Helsinki. Participants provided verbal and written informed consent.

## Supplementary Figure Legends

**Supplementary Figure 1.**
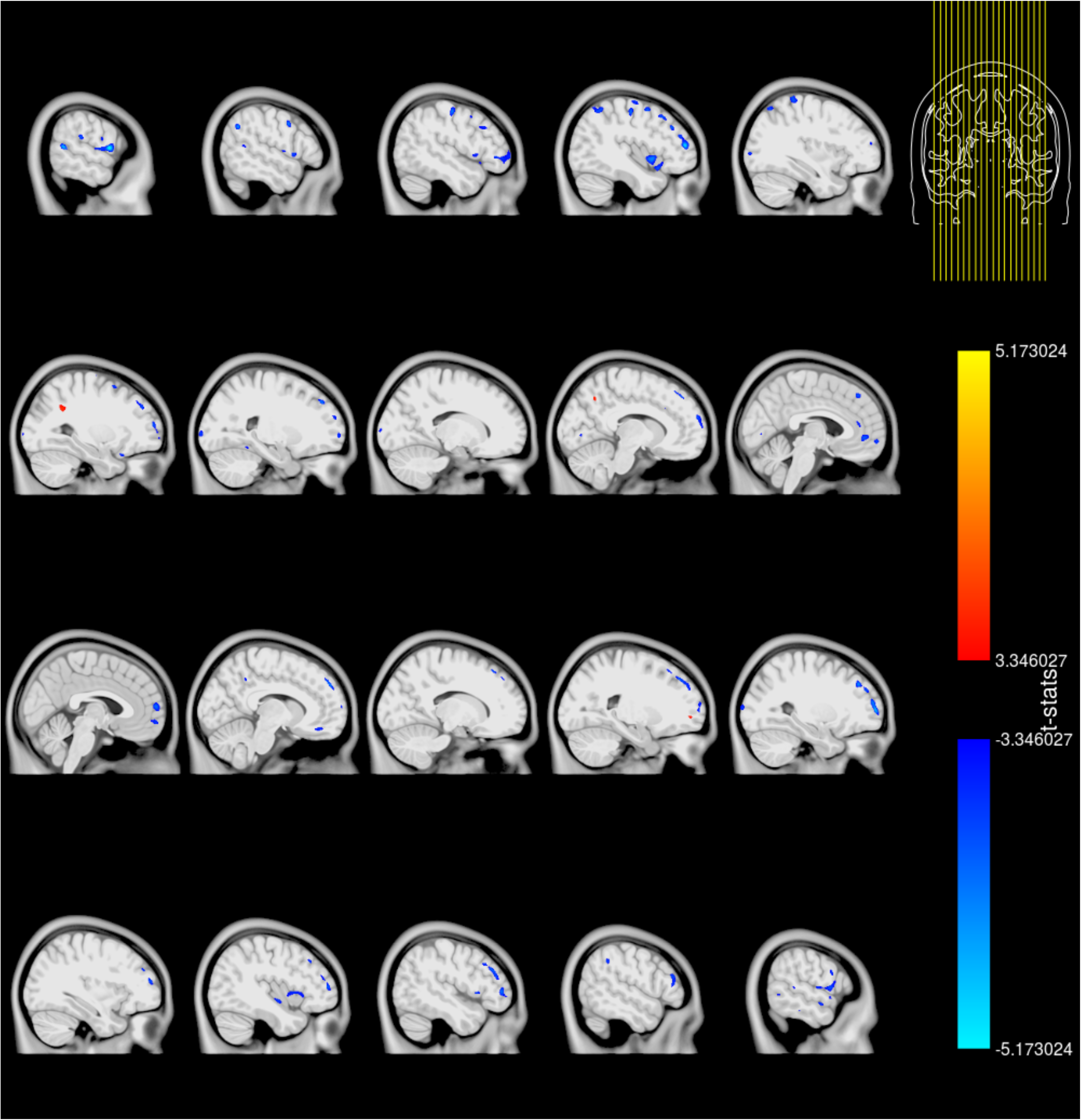
Voxel-based morphometry group contrast results (CA > HC).

**Supplementary Figure 2.**
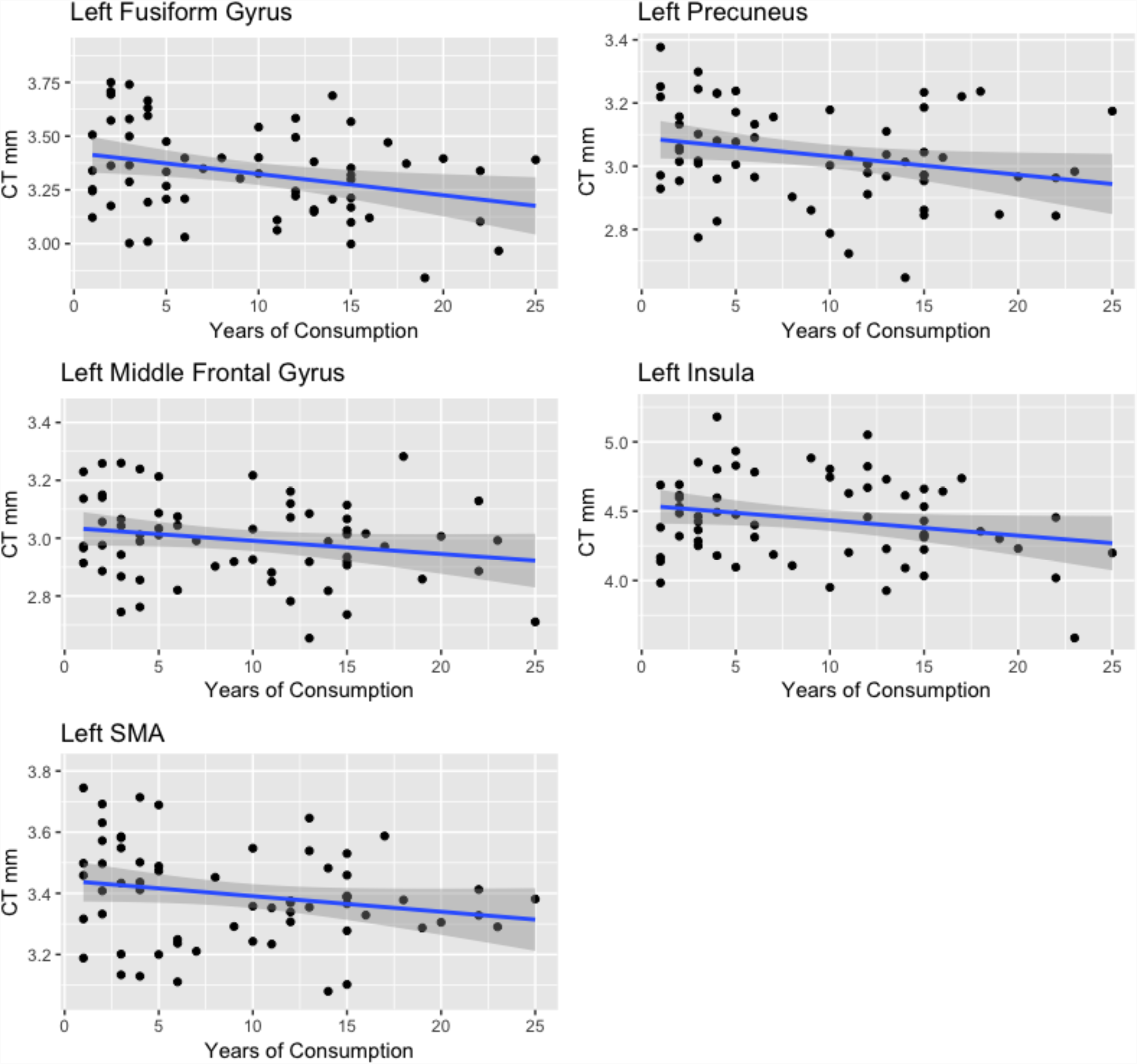
Scatterplots between CT peaks and years of consumption. CT = Cortical thickness.

**Supplementary Figure 3.**
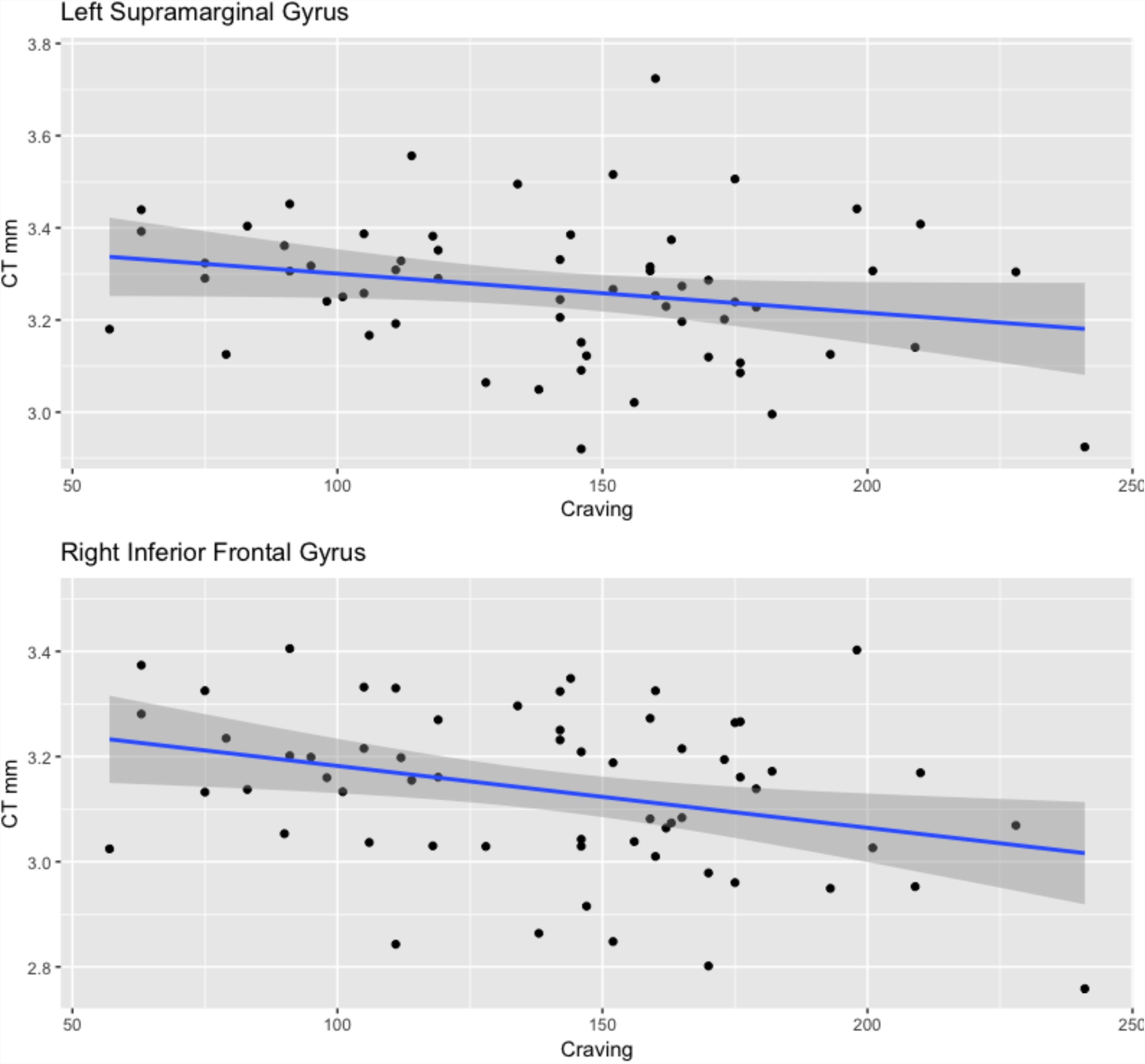
Scatterplots between CT peaks and cocaine craving. CT = Cortical thickness.

**Supplementary Figure 4.**
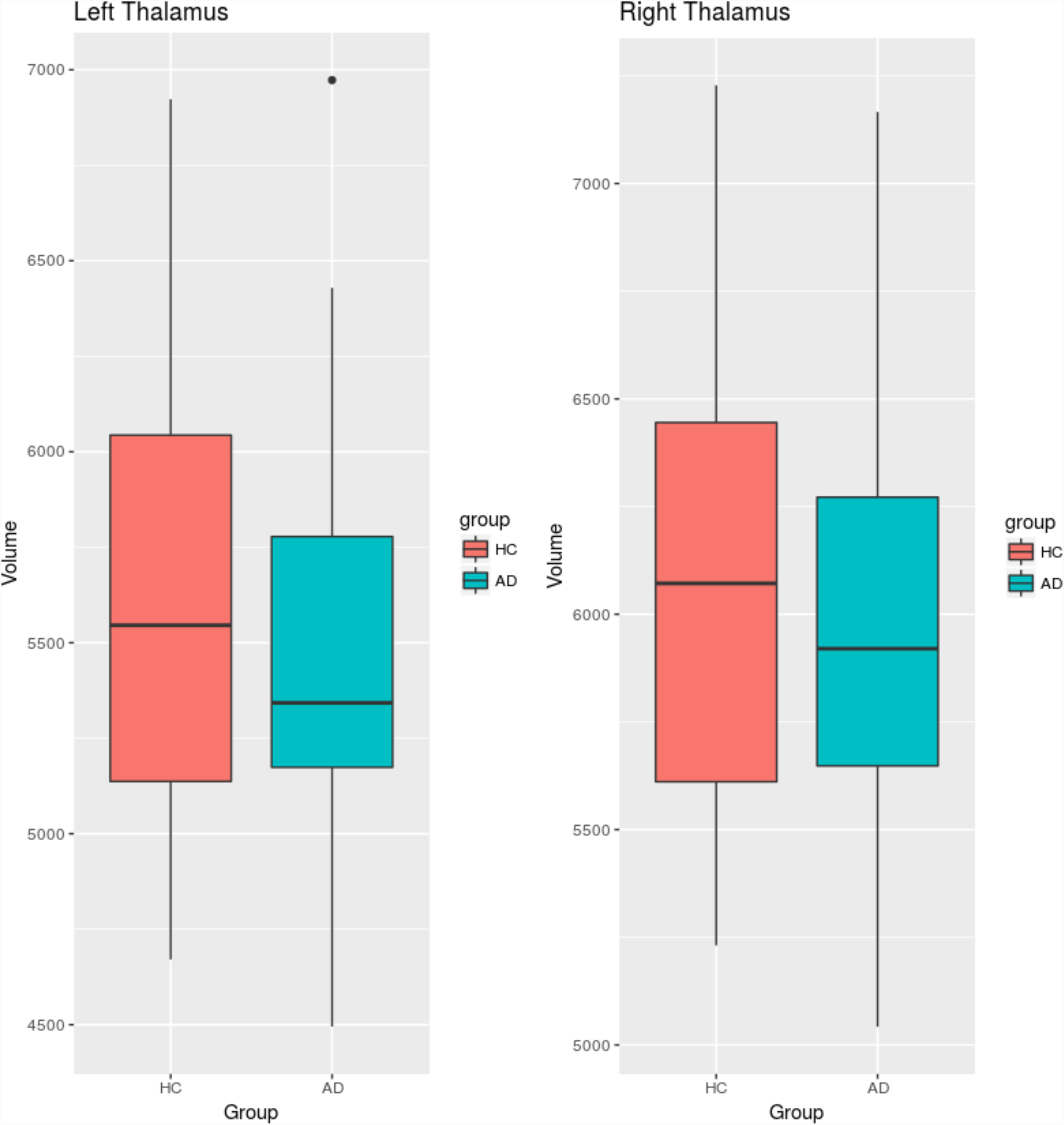
Boxplot of thalamus volume. HC = Healthy controls, AD = Cocaine addicts.

**Supplementary Figure 5.**
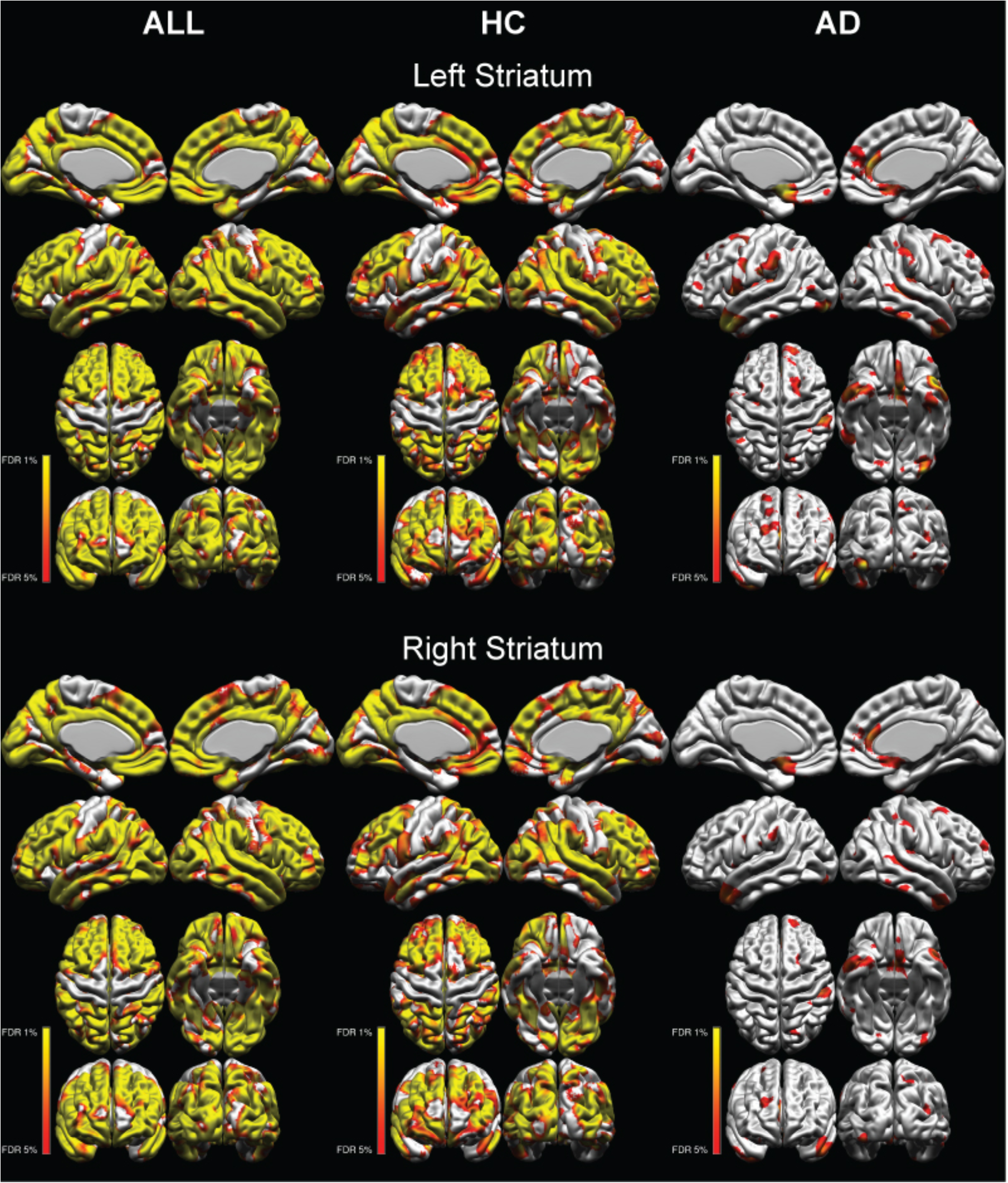
Correlation of left striatum volume and right striatum volume with cortical thickness. HC = Healthy controls, AD = Cocaine addicts, FDR = False Discovery Rate.

**Supplementary Table 1.** Voxel-based morphometry significant peaks.

| Hem | Brain | Area |  |  |  | x | y | z | t |
| --- | --- | --- | --- | --- | --- | --- | --- | --- | --- |
| Left | Precentral | Gyrus |  |  |  | -60 | 5 | 9 | -5.914 |
| Left | Frontal | Pole |  |  |  | -43 | 47 | 12 | -5.106 |
| Left | Precentral | Gyrus |  |  |  | -45 | -16 | 55 | -4.813 |
| Left | Middle | Temporal | Gyrus | temporooccipital | part | -61 | -51 | 11 | -4.763 |
| Left | Insular | Cortex |  |  |  | -44 | 8 | -6 | -4.709 |
| Left | Middle | Frontal | Gyrus |  |  | -44 | 31 | 33 | -4.509 |
| Left | Paracingulate | Gyrus |  |  |  | -7 | 24 | 32 | -4.507 |
| Left | Frontal | Pole |  |  |  | -50 | 37 | -3 | -4.469 |
| Left | Angular | Gyrus |  |  |  | -54 | -57 | 35 | -4.439 |
| Left | Frontal | Pole |  |  |  | -8 | 62 | 21 | -4.435 |
| Left | Superior | Frontal | Gyrus |  |  | -27 | 11 | 59 | -4.385 |
| Left | Precentral | Gyrus |  |  |  | -54 | 4 | 38 | -4.345 |
| Left | Middle | Frontal | Gyrus |  |  | -39 | 24 | 47 | -4.322 |
| Left | Frontal | Pole |  |  |  | -23 | 44 | 40 | -4.319 |
| Left | Frontal | Pole |  |  |  | -27 | 59 | 12 | -4.252 |
| Left | Temporal | Pole |  |  |  | -45 | 15 | -11 | -4.228 |
| Left | Superior | Frontal | Gyrus |  |  | -6 | 40 | 48 | -4.217 |
| Left | Angular | Gyrus |  |  |  | -43 | -56 | 53 | -4.174 |
| Left | Frontal | Pole |  |  |  | -9 | 43 | 47 | -4.139 |
| Left | Supramarginal | Gyrus |  | anterior | division | -62 | -30 | 23 | -4.122 |
| Left | Middle | Frontal | Gyrus |  |  | -42 | 2 | 55 | -4.098 |
| Left | Frontal | Pole |  |  |  | -28 | 44 | 35 | -4.065 |
| Left | Middle | Frontal | Gyrus |  |  | -43 | 18 | 47 | -4.047 |
| Left | Frontal | Pole |  |  |  | -5 | 63 | -7 | -4.031 |
| Left | Frontal | Pole |  |  |  | -49 | 43 | -4 | -4.026 |
| Left | Paracingulate | Gyrus |  |  |  | -4 | 47 | -3 | -4.025 |
| Left | Superior | Parietal | Lobule |  |  | -35 | -39 | 67 | -4.021 |
| Left | Frontal | Pole |  |  |  | -41 | 42 | 24 | -3.999 |
| Left | Frontal | Pole |  |  |  | -24 | 64 | 0 | -3.985 |
| Left | Lateral | Occipital | Cortex | inferior | division | -36 | -90 | 3 | -3.978 |
| Left | Lateral | Occipital | Cortex | superior | division | -35 | -67 | 56 | -3.889 |
| Left | Occipital | Pole |  |  |  | -24 | -102 | 1 | -3.877 |
| Left | Middle | Frontal | Gyrus |  |  | -47 | 24 | 35 | -3.84 |
| Left | Superior | Frontal | Gyrus |  |  | -10 | 38 | 51 | -3.819 |
| Left | Lateral | Occipital | Cortex | superior | division | -39 | -63 | 58 | -3.808 |
| Left | Frontal | Orbital | Cortex |  |  | -29 | 20 | -24 | -3.777 |
| Left | Frontal | Pole |  |  |  | -9 | 68 | 10 | -3.755 |
| Left | Frontal | Pole |  |  |  | -23 | 59 | 21 | -3.712 |
| Left | Frontal | Pole |  |  |  | -6 | 56 | 34 | -3.708 |
| Left | Precentral | Gyrus |  |  |  | -61 | -4 | 20 | -3.701 |
| Left | Supramarginal | Gyrus | anterior | division |  | -59 | -38 | 42 | -3.658 |
| Left | Frontal | Pole |  |  |  | -7 | 59 | 13 | -3.647 |
| Left | Temporal | Occipital | Fusiform |  |  | -24 | -47 | -15 | -3.643 |
| Left | Lingual | Gyrus |  |  |  | -9 | -78 | 0 | -3.637 |
| Left | Frontal | Pole |  |  |  | -48 | 53 | -1 | -3.636 |
| Left | Occipital | Pole |  |  |  | -15 | -103 | 6 | -3.624 |
| Left | Cingulate | Gyrus | anterior | division |  | -4 | 40 | 13 | -3.623 |
| Left | Superior | Frontal | Gyrus |  |  | -6 | 50 | 28 | -3.618 |
| Left | Middle | Frontal | Gyrus |  |  | -47 | 7 | 48 | -3.612 |
| Left | Frontal | Pole |  |  |  | -5 | 59 | 4 | -3.562 |
| Left | Cingulate | Gyrus | anterior | division |  | -5 | 34 | 23 | -3.516 |
| Left | Precentral | Gyrus |  |  |  | -31 | -12 | 72 | -3.512 |
| Left | Supramarginal | Gyrus | anterior | division |  | -51 | -33 | 34 | -3.491 |
| Left | Cingulate | Gyrus | anterior | division |  | -5 | 35 | 21 | -3.48 |
| Left | Inferior | Temporal | Gyrus | temporooccipital | part | -46 | -51 | -22 | -3.418 |
| Left | Occipital | Pole |  |  |  | -26 | -95 | -13 | -3.395 |
| Left | Postcentral | Gyrus |  |  |  | -47 | -29 | 62 | -3.381 |
| Left | Frontal | Orbital | Cortex |  |  | -38 | 23 | -15 | -3.363 |
| Left | Precuneus | Cortex |  |  |  | -11 | -61 | 44 | 3.808 |
| Right | Frontal | Pole |  |  |  | 27 | 61 | 5 | -4.865 |
| Right | Frontal | Pole |  |  |  | 37 | 54 | 12 | -4.577 |
| Right | Inferior | Frontal | Gyrus | pars | opercularis | 56 | 11 | 9 | -4.537 |
| Right | Frontal | Pole |  |  |  | 5 | 62 | 8 | -4.513 |
| Right | Superior | Frontal | Gyrus |  |  | 18 | 32 | 52 | -4.482 |
| Right | Frontal | Pole |  |  |  | 9 | 54 | 36 | -4.447 |
| Right | Superior | Frontal | Gyrus |  |  | 19 | 30 | 53 | -4.441 |
| Right | Frontal | Pole |  |  |  | 30 | 56 | 15 | -4.435 |
| Right | Frontal | Pole |  |  |  | 28 | 40 | 37 | -4.417 |
| Right | Frontal | Pole |  |  |  | 25 | 38 | 44 | -4.37 |
| Right | Inferior | Frontal | Gyrus | pars | opercularis | 57 | 13 | 16 | -4.345 |
| Right | Frontal | Pole |  |  |  | 46 | 45 | 3 | -4.337 |
| Right | Middle | Frontal | Gyrus |  |  | 48 | 31 | 29 | -4.248 |
| Right | Frontal | Pole |  |  |  | 31 | 46 | 31 | -4.235 |
| Right | Frontal | Pole |  |  |  | 45 | 38 | 23 | -4.186 |
| Right | Frontal | Pole |  |  |  | 10 | 50 | 41 | -4.169 |
| Right | Frontal | Pole |  |  |  | 26 | 50 | 29 | -4.139 |
| Right | Middle | Frontal | Gyrus |  |  | 51 | 32 | 23 | -4.129 |
| Right | Supramarginal | Gyrus |  | anterior | division | 63 | -21 | 30 | -4.064 |
| Right | Middle | Frontal | Gyrus |  |  | 40 | 30 | 40 | -4.049 |
| Right | Angular | Gyrus |  |  |  | 51 | -45 | 39 | -4.023 |
| Right | Frontal | Pole |  |  |  | 20 | 50 | 37 | -3.993 |
| Right | Frontal | Pole |  |  |  | 4 | 61 | -10 | -3.991 |
| Right | Precentral | Gyrus |  |  |  | 59 | 6 | 8 | -3.976 |
| Right | Frontal | Pole |  |  |  | 21 | 47 | 39 | -3.957 |
| Right | Frontal | Medial | Cortex |  |  | 11 | 41 | -16 | -3.944 |
| Right | Middle | Frontal | Gyrus |  |  | 46 | 24 | 37 | -3.937 |
| Right | Frontal | Operculum | Cortex |  |  | 44 | 14 | 2 | -3.934 |
| Right | Occipital | Pole |  |  |  | 27 | -100 | 8 | -3.883 |
| Right | Precentral | Gyrus |  |  |  | 60 | 7 | 25 | -3.882 |
| Right | Central | Opercular | Cortex |  |  | 58 | -8 | 8 | -3.873 |
| Right | Frontal | Pole |  |  |  | 45 | 49 | -2 | -3.864 |
| Right | Middle | Temporal | Gyrus | anterior | division | 60 | -5 | -12 | -3.843 |
| Right | Planum | Polare |  |  |  | 42 | -9 | -8 | -3.825 |
| Right | Superior | Temporal | Gyrus | posterior | division | 64 | -36 | 8 | -3.802 |
| Right | Precuneous | Cortex |  |  |  | 7 | -48 | 43 | -3.8 |
| Right | Frontal | Pole |  |  |  | 0 | 60 | -9 | -3.776 |
| Right | Inferior | Frontal | Gyrus | pars | triangularis | 54 | 32 | 11 | -3.762 |
| Right | Inferior | Frontal | Gyrus | pars | triangularis | 53 | 34 | 14 | -3.757 |
| Right | Frontal | Pole |  |  |  | 6 | 58 | 18 | -3.73 |
| Right | Frontal | Pole |  |  |  | 23 | 65 | 13 | -3.722 |
| Right | Inferior | Frontal | Gyrus | pars | opercularis | 56 | 21 | 10 | -3.643 |
| Right | Frontal | Operculum | Cortex |  |  | 41 | 21 | -2 | -3.587 |
| Right | Occipital | Fusiform | Gyrus |  |  | 20 | -72 | -11 | -3.578 |
| Right | Middle | Temporal | Gyrus | posterior | division | 69 | -10 | -19 | -3.571 |
| Right | Middle | Temporal | Gyrus | temporooccipital | part | 60 | -56 | -1 | -3.56 |
| Right | Postcentral | Gyrus |  |  |  | 43 | -21 | 51 | -3.504 |
| Right | Precentral | Gyrus |  |  |  | 56 | -1 | 40 | -3.494 |
| Right | Superior | Temporal | Gyrus | anterior | division | 59 | 6 | -4 | -3.492 |
| Right | Inferior | Temporal | Gyrus | posterior | division | 59 | -31 | -20 | -3.43 |
| Right | Postcentral | Gyrus |  |  |  | 43 | -22 | 65 | -3.413 |
| Right | Lateral | Occipital | Cortex | inferior | division | 39 | -90 | -4 | -3.403 |
| Right | Frontal | Pole |  |  |  | 45 | 56 | -13 | -3.374 |
| Right | Precentral | Gyrus |  |  |  | 4 | -21 | 54 | -3.349 |
| Right | Temporal | Pole |  |  |  | 42 | 11 | -16 | -3.347 |
| Right | Frontal | Pole |  |  |  | 21 | 54 | -3 | 3.644 |

Hem = Hemisphere, t = t-value, coordinates are in MNI.

**Supplementary Table 2.**
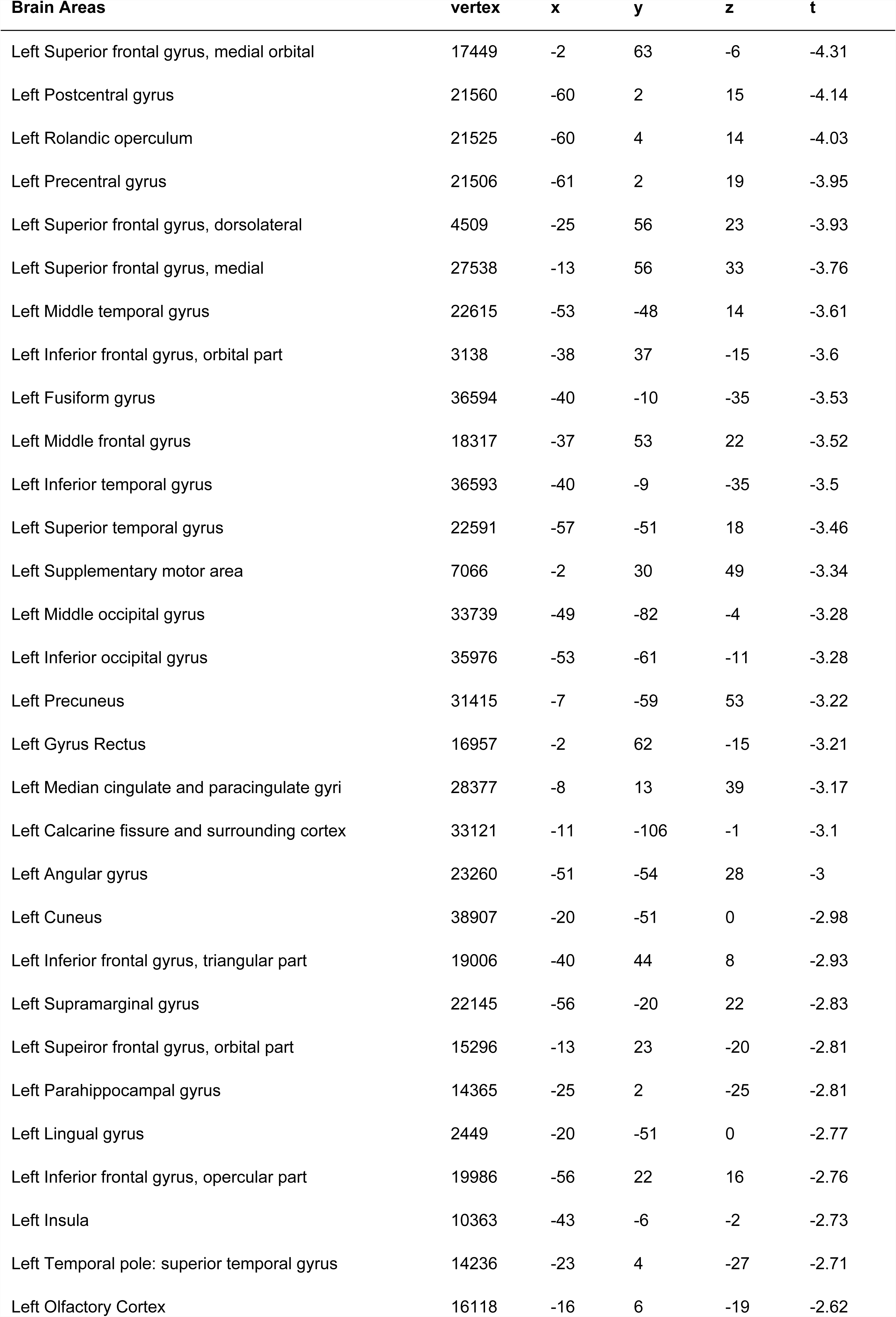
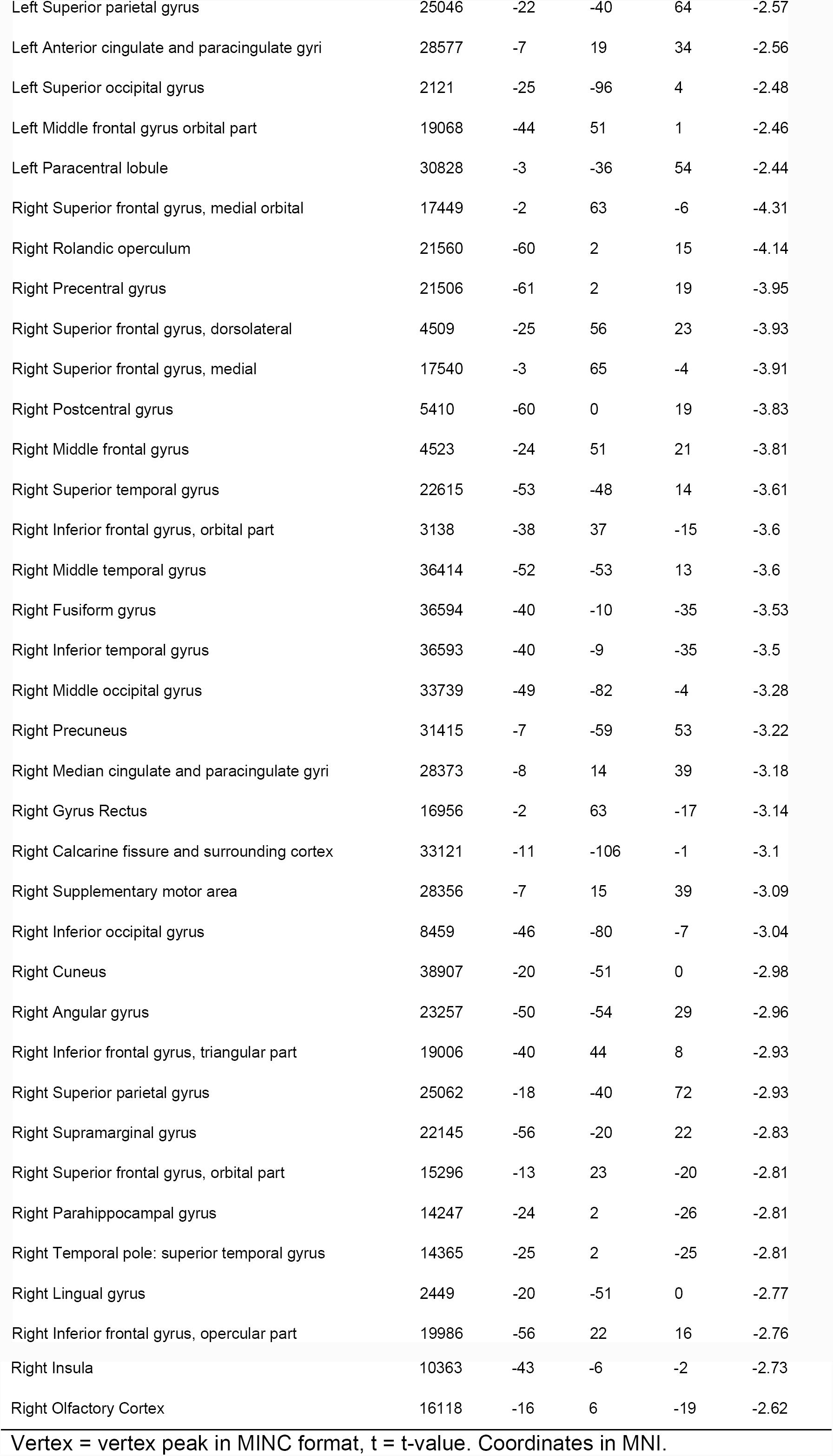
Cortical thickness significant peaks at FDR 10%.

| Brain Areas | vertex | x | y | z | t |
| --- | --- | --- | --- | --- | --- |
| Left Superior frontal gyrus, medial orbital | 17449 | -2 | 63 | -6 | -4.31 |
| Left Postcentral gyrus | 21560 | -60 | 2 | 15 | -4.14 |
| Left Rolandic operculum | 21525 | -60 | 4 | 14 | -4.03 |
| Left Precentral gyrus | 21506 | -61 | 2 | 19 | -3.95 |
| Left Superior frontal gyrus, dorsolateral | 4509 | -25 | 56 | 23 | -3.93 |
| Left Superior frontal gyrus, medial | 27538 | -13 | 56 | 33 | -3.76 |
| Left Middle temporal gyrus | 22615 | -53 | -48 | 14 | -3.61 |
| Left Inferior frontal gyrus, orbital part | 3138 | -38 | 37 | -15 | -3.6 |
| Left Fusiform gyrus | 36594 | -40 | -10 | -35 | -3.53 |
| Left Middle frontal gyrus | 18317 | -37 | 53 | 22 | -3.52 |
| Left Inferior temporal gyrus | 36593 | -40 | -9 | -35 | -3.5 |
| Left Superior temporal gyrus | 22591 | -57 | -51 | 18 | -3.46 |
| Left Supplementary motor area | 7066 | -2 | 30 | 49 | -3.34 |
| Left Middle occipital gyrus | 33739 | -49 | -82 | -4 | -3.28 |
| Left Inferior occipital gyrus | 35976 | -53 | -61 | -11 | -3.28 |
| Left Precuneus | 31415 | -7 | -59 | 53 | -3.22 |
| Left Gyrus Rectus | 16957 | -2 | 62 | -15 | -3.21 |
| Left Median cingulate and paracingulate gyri | 28377 | -8 | 13 | 39 | -3.17 |
| Left Calcarine fissure and surrounding cortex | 33121 | -11 | -106 | -1 | -3.1 |
| Left Angular gyrus | 23260 | -51 | -54 | 28 | -3 |
| Left Cuneus | 38907 | -20 | -51 | 0 | -2.98 |
| Left Inferior frontal gyrus, triangular part | 19006 | -40 | 44 | 8 | -2.93 |
| Left Supramarginal gyrus | 22145 | -56 | -20 | 22 | -2.83 |
| Left Superior frontal gyrus, orbital part | 15296 | -13 | 23 | -20 | -2.81 |
| Left Parahippocampal gyrus | 14365 | -25 | 2 | -25 | -2.81 |
| Left Lingual gyrus | 2449 | -20 | -51 | 0 | -2.77 |
| Left Inferior frontal gyrus, opercular part | 19986 | -56 | 22 | 16 | -2.76 |
| Left Insula | 10363 | -43 | -6 | -2 | -2.73 |
| Left Temporal pole: superior temporal gyrus | 14236 | -23 | 4 | -27 | -2.71 |
| Left Olfactory Cortex | 16118 | -16 | 6 | -19 | -2.62 |
| Left Superior parietal gyrus | 25046 | -22 | -40 | 64 | -2.57 |
| Left Anterior cingulate and paracingulate gyri | 28577 | -7 | 19 | 34 | -2.56 |
| Left Superior occipital gyrus | 2121 | -25 | -96 | 4 | -2.48 |
| Left Middle frontal gyrus orbital part | 19068 | -44 | 51 | 1 | -2.46 |
| Left Paracentral lobule | 30828 | -3 | -36 | 54 | -2.44 |
| Right Superior frontal gyrus, medial orbital | 17449 | -2 | 63 | -6 | -4.31 |
| Right Rolandic operculum | 21560 | -60 | 2 | 15 | -4.14 |
| Right Precentral gyrus | 21506 | -61 | 2 | 19 | -3.95 |
| Right Superior frontal gyrus, dorsolateral | 4509 | -25 | 56 | 23 | -3.93 |
| Right Superior frontal gyrus, medial | 17540 | -3 | 65 | -4 | -3.91 |
| Right Postcentral gyrus | 5410 | -60 | 0 | 19 | -3.83 |
| Right Middle frontal gyrus | 4523 | -24 | 51 | 21 | -3.81 |
| Right Superior temporal gyrus | 22615 | -53 | -48 | 14 | -3.61 |
| Right Inferior frontal gyrus, orbital part | 3138 | -38 | 37 | -15 | -3.6 |
| Right Middle temporal gyrus | 36414 | -52 | -53 | 13 | -3.6 |
| Right Fusiform gyrus | 36594 | -40 | -10 | -35 | -3.53 |
| Right Inferior temporal gyrus | 36593 | -40 | -9 | -35 | -3.5 |
| Right Middle occipital gyrus | 33739 | -49 | -82 | -4 | -3.28 |
| Right Precuneus | 31415 | -7 | -59 | 53 | -3.22 |
| Right Median cingulate and paracingulate gyri | 28373 | -8 | 14 | 39 | -3.18 |
| Right Gyrus Rectus | 16956 | -2 | 63 | -17 | -3.14 |
| Right Calcarine fissure and surrounding cortex | 33121 | -11 | -106 | -1 | -3.1 |
| Right Supplementary motor area | 28356 | -7 | 15 | 39 | -3.09 |
| Right Inferior occipital gyrus | 8459 | -46 | -80 | -7 | -3.04 |
| Right Cuneus | 38907 | -20 | -51 | 0 | -2.98 |
| Right Angular gyrus | 23257 | -50 | -54 | 29 | -2.96 |
| Right Inferior frontal gyrus, triangular part | 19006 | -40 | 44 | 8 | -2.93 |
| Right Superior parietal gyrus | 25062 | -18 | -40 | 72 | -2.93 |
| Right Supramarginal gyrus | 22145 | -56 | -20 | 22 | -2.83 |
| Right Superior frontal gyrus, orbital part | 15296 | -13 | 23 | -20 | -2.81 |
| Right Parahippocampal gyrus | 14247 | -24 | 2 | -26 | -2.81 |
| Right Temporal pole: superior temporal gyrus | 14365 | -25 | 2 | -25 | -2.81 |
| Right Lingual gyrus | 2449 | -20 | -51 | 0 | -2.77 |
| Right Inferior frontal gyrus, opercular part | 19986 | -56 | 22 | 16 | -2.76 |
| Right Insula | 10363 | -43 | -6 | -2 | -2.73 |
| Right Olfactory Cortex | 16118 | -16 | 6 | -19 | -2.62 |

Vertex = vertex peak in MINC format, t = t-value. Coordinates in MNI.

**Supplementary Table 3.** Correlations between CT peaks and years of cocaine use.

| Brain Area | vertex | r | uncorr-p | fdr-p |
| --- | --- | --- | --- | --- |
| Left Fusiform Gyrus | 36594 | -0.35 | 0.003 | 0.02 |
| Left Middle Frontal Gyrus | 18317 | -0.20 | 0.057 | 0.057 |
| Left SMA | 7066 | -0.24 | 0.031 | 0.05 |
| Left Precuneus | 31415 | -0.22 | 0.047 | 0.057 |
| Left Insula | 10363 | -0.26 | 0.023 | 0.046 |

Vertex = Surface vertex CIVET 1.1.12, r = correlation coefficient, uncorr-p = non-corrected p value, fdr-p = p value corrected for multiple comparisons using false discovery rate.

**Supplementary Table 4.** Correlations between CT peaks and cocaine craving.

| Brain Area | vertex | r | uncorr-p | fdr-p |
| --- | --- | --- | --- | --- |
| Left Supramarginal Gyrus | 22145 | -0.24 | 0.034 | 0.045 |
| Right Middle Frontal Gyrus | 18285 | -0.22 | 0.042 | 0.045 |
| Right Lingual Gyrus | 9552 | -0.22 | 0.045 | 0.045 |
| Right Parahippocampal Gyrus | 38137 | -0.24 | 0.033 | 0.045 |

Vertex = Surface vertex CIVET 1.1.12, r = correlation coefficient, uncorr-p = non-corrected p value, fdr-p = p value corrected for multiple comparisons using false discovery rate.

**Supplementary Table 5.** Left Striatum volume significant peaks of correlation with cortical thickness in ALL participants.

| Brain Area | vertex | x | y | z | t |
| --- | --- | --- | --- | --- | --- |
| Left Calcarine fissure and surrounding cortex | 33295 | -11 | -92 | -19 | 6 |
| Left Olfactory Cortex | 4040 | -2 | 17 | -14 | 5.96 |
| Left Lingual gyrus | 39387 | -12 | -87 | -20 | 5.91 |
| Left Middle temporal gyrus | 13331 | -53 | 12 | -30 | 5.61 |
| Left Anterior cingulate and paracingulate gyri | 16051 | -3 | 20 | -13 | 5.59 |
| Left Superior temporal gyrus | 22384 | -45 | -34 | 17 | 5.5 |
| Left Gyrus Rectus | 15115 | -6 | 39 | -28 | 5.39 |
| Left Inferior occipital gyrus | 33480 | -34 | -90 | -10 | 5.3 |
| Left Precuneus | 40676 | -13 | -52 | 2 | 5.3 |
| Left Superior frontal gyrus, medial orbital | 15921 | -5 | 25 | -14 | 5.29 |
| Left Middle occipital gyrus | 33446 | -33 | -93 | -10 | 5.23 |
| Left Supramarginal gyrus | 22244 | -43 | -33 | 16 | 5.11 |
| Left Rolandic operculum | 5493 | -37 | -26 | 19 | 5.07 |
| Left Fusiform gyrus | 35422 | -40 | -75 | -16 | 5.01 |
| Left Heschl gyrus | 672 | -39 | -31 | 18 | 5.01 |
| Left Temporal pole: superior temporal gyrus | 13189 | -50 | 16 | -29 | 5 |
| Left Temporal pole: middle temporal gyrus | 3348 | -49 | 14 | -32 | 4.99 |
| Left Insula | 10525 | -39 | -17 | 5 | 4.97 |
| Left Precentral gyrus | 20887 | -53 | -5 | 30 | 4.96 |
| Left Inferior frontal gyrus, opercular part | 20608 | -40 | 12 | 28 | 4.96 |
| Left Angular gyrus | 23649 | -48 | -48 | 49 | 4.96 |
| Left Middle frontal gyrus | 20597 | -39 | 11 | 28 | 4.94 |
| Left Inferior frontal gyrus, triangular part | 20611 | -41 | 12 | 28 | 4.93 |
| Left Cuneus | 38388 | -15 | -55 | 4 | 4.91 |
| Left Superior frontal gyrus, dorsolateral | 6720 | -23 | -5 | 55 | 4.89 |
| Left Inferior parietal, but supramarginal and angular gyri | 23531 | -47 | -47 | 47 | 4.75 |
| Left Median cingulate and paracingulate gyri | 7483 | -5 | -18 | 44 | 4.66 |
| Left Postcentral gyrus | 20865 | -53 | -6 | 29 | 4.66 |
| Left Supplementary motor area | 1772 | -8 | 3 | 45 | 4.54 |
| Left Superior frontal gyrus, medial | 28049 | -4 | 42 | 35 | 4.32 |
| Left Posterior cingulate gyrus | 40909 | -2 | -52 | 16 | 4.31 |
| Left Superior occipital gyrus | 33893 | -27 | -85 | 21 | 4.28 |
| Left Inferior temporal gyrus | 36703 | -45 | -24 | -26 | 4.17 |
| Left Superior parietal gyrus | 25063 | -29 | -40 | 59 | 4.13 |
| Left Middle frontal gyrus orbital part | 19079 | -42 | 52 | 1 | 4.1 |
| Left Superior frontal gyrus, orbital part | 15206 | -10 | 40 | -23 | 3.88 |
| Left Paracentral lobule | 30979 | -13 | -44 | 62 | 3.79 |
| Left Parahippocampal gyrus | 37578 | -30 | -50 | -8 | 3.57 |
| Left Inferior frontal gyrus, orbital part | 18643 | -39 | 52 | -13 | 3.45 |
| Right Rolandic operculum | 21843 | -40 | -24 | 18 | 6.22 |
| Right Superior temporal gyrus | 13008 | -45 | -2 | -11 | 6.02 |
| Right Anterior cingulate and paracingulate gyri | 16045 | -3 | 16 | -14 | 5.99 |
| Right Supramarginal gyrus | 21832 | -42 | -25 | 19 | 5.97 |
| Right Inferior frontal gyrus, triangular part | 19447 | -42 | 22 | 7 | 5.8 |
| Right Olfactory Cortex | 16042 | -3 | 15 | -15 | 5.79 |
| Right Heschl gyrus | 10642 | -40 | -30 | 15 | 5.79 |
| Right Insula | 10569 | -36 | -23 | 16 | 5.73 |
| Right Middle occipital gyrus | 34433 | -41 | -88 | 13 | 5.65 |
| Right Median cingulate and paracingulate gyri | 29837 | -11 | -21 | 41 | 5.58 |
| Right Postcentral gyrus | 21790 | -47 | -16 | 15 | 5.52 |
| Right Temporal pole: superior temporal gyrus | 3279 | -45 | 5 | -14 | 5.47 |
| Right Superior frontal gyrus, medial | 4440 | -4 | 52 | 25 | 5.4 |
| Right Inferior frontal gyrus, opercular part | 19526 | -42 | 20 | 7 | 5.39 |
| Right Middle temporal gyrus | 11030 | -55 | -18 | -8 | 5.36 |
| Right Superior occipital gyrus | 34019 | -26 | -84 | 18 | 5.26 |
| Right Posterior cingulate gyrus | 30208 | -7 | -51 | 32 | 5.21 |
| Right Inferior parietal, but supramarginal and angular gyri | 22744 | -49 | -31 | 44 | 5.18 |
| Right Temporal pole: middle temporal gyrus | 3374 | -53 | 9 | -39 | 5.18 |
| Right Lingual gyrus | 39369 | -15 | -82 | -18 | 5.14 |
| Right Precuneus | 7934 | -10 | -76 | 50 | 5.14 |
| Right Superior frontal gyrus, orbital part | 888 | -24 | 29 | -20 | 5.06 |
| Right Superior parietal gyrus | 31649 | -11 | -74 | 50 | 5.01 |
| Right Supplementary motor area | 7030 | -9 | -1 | 46 | 4.94 |
| Right Middle frontal gyrus | 19788 | -33 | 38 | 31 | 4.9 |
| Right Superior frontal gyrus, medial orbital | 4028 | -2 | 20 | -17 | 4.84 |
| Right Inferior temporal gyrus | 856 | -47 | 2 | -41 | 4.8 |
| Right Inferior frontal gyrus, orbital part | 13926 | -24 | 28 | -18 | 4.71 |
| Right Precentral gyrus | 20657 | -44 | 9 | 24 | 4.66 |
| Right Superior frontal gyrus, dorsolateral | 27336 | -18 | 54 | 37 | 4.62 |
| Right Cuneus | 31915 | -17 | -67 | 20 | 4.33 |
| Right Angular gyrus | 24041 | -28 | -67 | 33 | 4.32 |
| Right Calcarine fissure and surrounding cortex | 39388 | -9 | -86 | -18 | 4.24 |
| Right Fusiform gyrus | 37897 | -25 | -76 | -13 | 4.21 |
| Right Gyrus Rectus | 3841 | -11 | 28 | -23 | 4.2 |
| Right Paracentral lobule | 1862 | -9 | -26 | 46 | 3.98 |
| Right Middle frontal gyrus orbital part | 18535 | -23 | 54 | -15 | 3.79 |
| Right Inferior occipital gyrus | 33607 | -35 | -91 | -7 | 3.78 |
| Right Parahippocampal gyrus | 14247 | -24 | 2 | -26 | 3.65 |

**Supplementary Table 6.** Right Striatum volume significant peaks of correlation with cortical thickness in ALL participants.

| Brain Area | vertex | x | y | z | t |
| --- | --- | --- | --- | --- | --- |
| Left Calcarine fissure and surrounding cortex | 33295 | -11 | -92 | -19 | 6.02 |
| Left Olfactory Cortex | 16045 | -3 | 16 | -14 | 5.9 |
| Left Lingual gyrus | 39387 | -12 | -87 | -20 | 5.84 |
| Left Anterior cingulate and paracingulate gyri | 16051 | -3 | 20 | -13 | 5.33 |
| Left Precuneus | 10176 | -12 | -52 | 3 | 5.29 |
| Left Superior frontal gyrus, medial orbital | 16931 | -4 | 57 | -14 | 5.25 |
| Left Middle temporal gyrus | 13331 | -53 | 12 | -30 | 5.22 |
| Left Gyrus Rectus | 16932 | -3 | 58 | -15 | 5.16 |
| Left Superior temporal gyrus | 22384 | -45 | -34 | 17 | 4.98 |
| Left Insula | 10525 | -39 | -17 | 5 | 4.96 |
| Left Middle occipital gyrus | 537 | -33 | -92 | -10 | 4.93 |
| Left Inferior occipital gyrus | 33480 | -34 | -90 | -10 | 4.93 |
| Left Rolandic operculum | 21836 | -40 | -25 | 19 | 4.86 |
| Left Supramarginal gyrus | 21837 | -40 | -26 | 20 | 4.85 |
| Left Cuneus | 9619 | -14 | -56 | 5 | 4.83 |
| Left Angular gyrus | 23649 | -48 | -48 | 49 | 4.8 |
| Left Fusiform gyrus | 35422 | -40 | -75 | -16 | 4.78 |
| Left Inferior frontal gyrus, triangular part | 11869 | -32 | 21 | 11 | 4.76 |
| Left Temporal pole: superior temporal gyrus | 13188 | -49 | 16 | -30 | 4.75 |
| Left Inferior frontal gyrus, opercular part | 20608 | -40 | 12 | 28 | 4.73 |
| Left Middle frontal gyrus | 20597 | -39 | 11 | 28 | 4.72 |
| Left Heschl gyrus | 2642 | -39 | -19 | 4 | 4.69 |
| Left Temporal pole: middle temporal gyrus | 3348 | -49 | 14 | -32 | 4.65 |
| Left Precentral gyrus | 26764 | -23 | -5 | 54 | 4.58 |
| Left Superior frontal gyrus, dorsolateral | 26745 | -23 | -4 | 54 | 4.58 |
| Left Inferior parietal, but supramarginal and angular gyri | 23531 | -47 | -47 | 47 | 4.56 |
| Left Median cingulate and paracingulate gyri | 29812 | -4 | -19 | 44 | 4.5 |
| Left Postcentral gyrus | 22201 | -48 | -23 | 19 | 4.48 |
| Left Posterior cingulate gyrus | 40909 | -2 | -52 | 16 | 4.46 |
| Left Superior occipital gyrus | 33893 | -27 | -85 | 21 | 4.37 |
| Left Inferior temporal gyrus | 36703 | -45 | -24 | -26 | 4.16 |
| Left Superior frontal gyrus, medial | 7043 | -5 | 43 | 34 | 4.1 |
| Left Supplementary motor area | 1772 | -8 | 3 | 45 | 4.06 |
| Left Superior parietal gyrus | 23435 | -32 | -55 | 43 | 3.97 |
| Left Middle frontal gyrus orbital part | 18981 | -33 | 58 | 3 | 3.78 |
| Left Superior frontal gyrus, orbital part | 15200 | -11 | 37 | -23 | 3.71 |
| Left Paracentral lobule | 30979 | -13 | -44 | 62 | 3.45 |
| Left Parahippocampal gyrus | 37578 | -30 | -50 | -8 | 3.4 |
| Left Inferior frontal gyrus, orbital part | 18643 | -39 | 52 | -13 | 3.19 |
| Right Anterior cingulate and paracingulate gyri | 16045 | -3 | 16 | -14 | 6.17 |
| Right Olfactory Cortex | 16042 | -3 | 15 | -15 | 5.99 |
| Right Rolandic operculum | 21843 | -40 | -24 | 18 | 5.98 |
| Right Middle occipital gyrus | 34433 | -41 | -88 | 13 | 5.86 |
| Right Supramarginal gyrus | 21837 | -40 | -26 | 20 | 5.77 |
| Right Superior temporal gyrus | 13008 | -45 | -2 | -11 | 5.68 |
| Right Median cingulate and paracingulate gyri | 29682 | -2 | -2 | 44 | 5.59 |
| Right Heschl gyrus | 10639 | -39 | -30 | 17 | 5.59 |
| Right Inferior frontal gyrus, triangular part | 19447 | -42 | 22 | 7 | 5.57 |
| Right Insula | 10569 | -36 | -23 | 16 | 5.47 |
| Right Superior occipital gyrus | 34019 | -26 | -84 | 18 | 5.28 |
| Right Inferior frontal gyrus, opercular part | 19526 | -42 | 20 | 7 | 5.19 |
| Right Postcentral gyrus | 21790 | -47 | -16 | 15 | 5.12 |
| Right Lingual gyrus | 39369 | -15 | -82 | -18 | 5.06 |
| Right Superior frontal gyrus, orbital part | 888 | -24 | 29 | -20 | 5.05 |
| Right Superior frontal gyrus, medial | 4440 | -4 | 52 | 25 | 5.04 |
| Right Middle temporal gyrus | 9114 | -54 | -50 | 11 | 5.03 |
| Right Temporal pole: superior temporal gyrus | 3279 | -45 | 5 | -14 | 4.99 |
| Right Inferior parietal, but supramarginal and angular gyri | 22744 | -49 | -31 | 44 | 4.97 |
| Right Supplementary motor area | 7030 | -9 | -1 | 46 | 4.95 |
| Right Precuneus | 7934 | -10 | -76 | 50 | 4.93 |
| Right Superior parietal gyrus | 31655 | -12 | -74 | 50 | 4.9 |
| Right Posterior cingulate gyrus | 30152 | -4 | -52 | 26 | 4.88 |
| Right Temporal pole: middle temporal gyrus | 3374 | -53 | 9 | -39 | 4.88 |
| Right Superior frontal gyrus, medial orbital | 15996 | -1 | 18 | -17 | 4.82 |
| Right Middle frontal gyrus | 19788 | -33 | 38 | 31 | 4.6 |
| Right Inferior frontal gyrus, orbital part | 13923 | -24 | 27 | -19 | 4.59 |
| Right Inferior temporal gyrus | 856 | -47 | 2 | -41 | 4.42 |
| Right Superior frontal gyrus, dorsolateral | 27336 | -18 | 54 | 37 | 4.39 |
| Right Precentral gyrus | 20651 | -44 | 9 | 25 | 4.36 |
| Right Cuneus | 31915 | -17 | -67 | 20 | 4.19 |
| Right Fusiform gyrus | 9422 | -35 | -54 | -19 | 4.16 |
| Right Angular gyrus | 24041 | -28 | -67 | 33 | 4.15 |
| Right Calcarine fissure and surrounding cortex | 39388 | -9 | -86 | -18 | 4.05 |
| Right Gyrus Rectus | 3841 | -11 | 28 | -23 | 4.02 |
| Right Paracentral lobule | 1862 | -9 | -26 | 46 | 3.91 |
| Right Inferior occipital gyrus | 33607 | -35 | -91 | -7 | 3.85 |
| Right Middle frontal gyrus orbital part | 18538 | -23 | 51 | -14 | 3.74 |
| Right Parahippocampal gyrus | 14247 | -24 | 2 | -26 | 3.55 |

**Supplementary Table 7.**
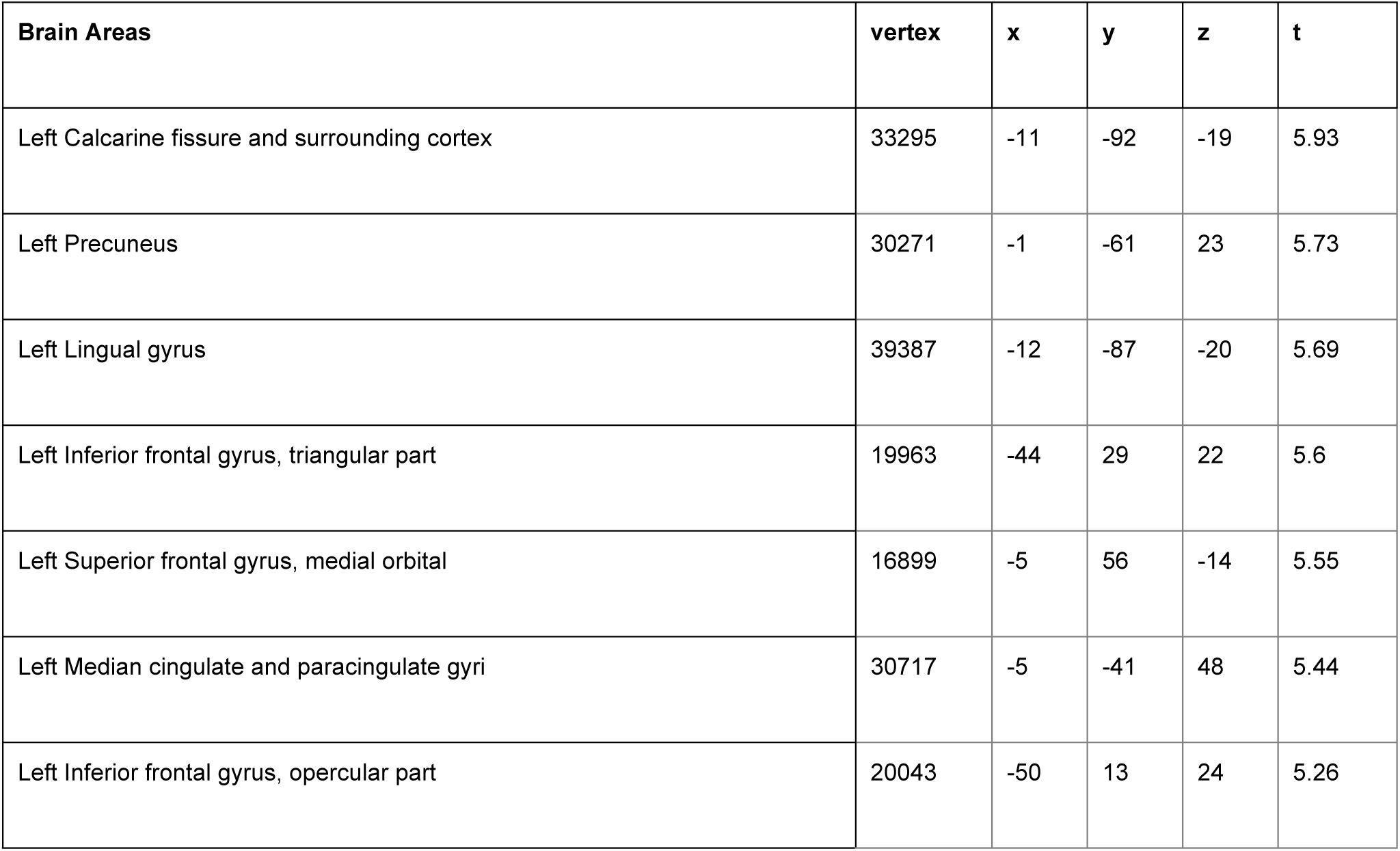
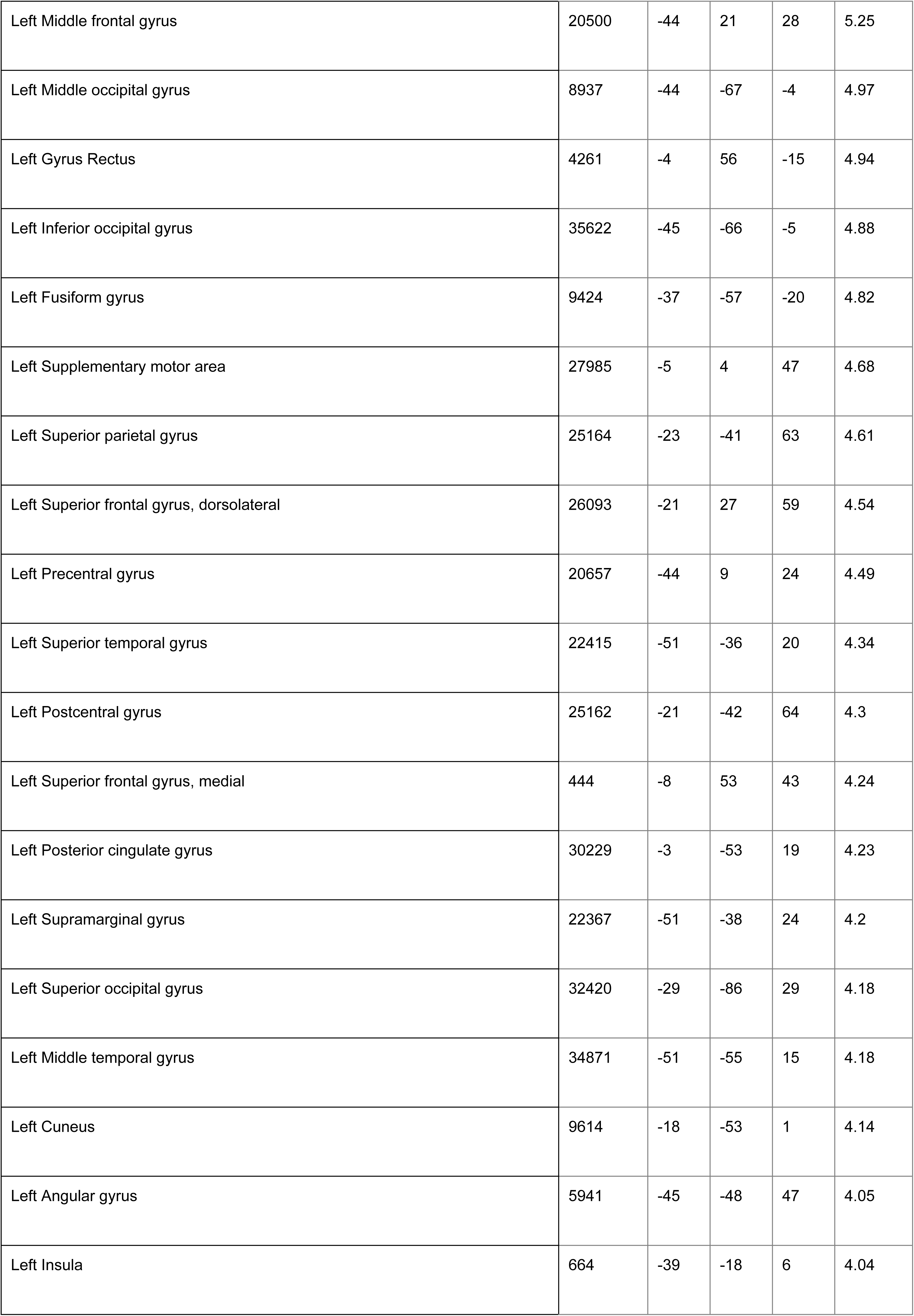
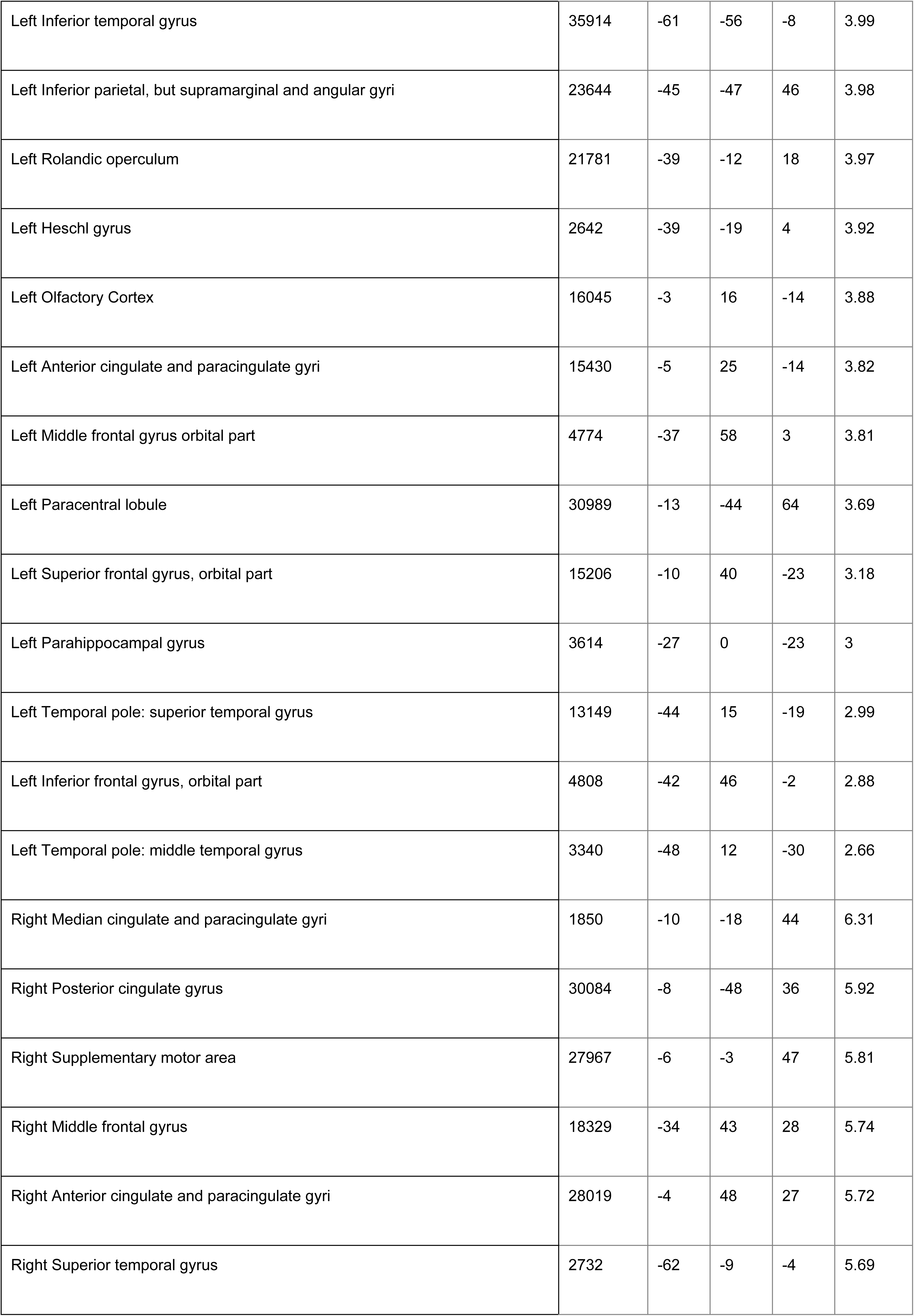
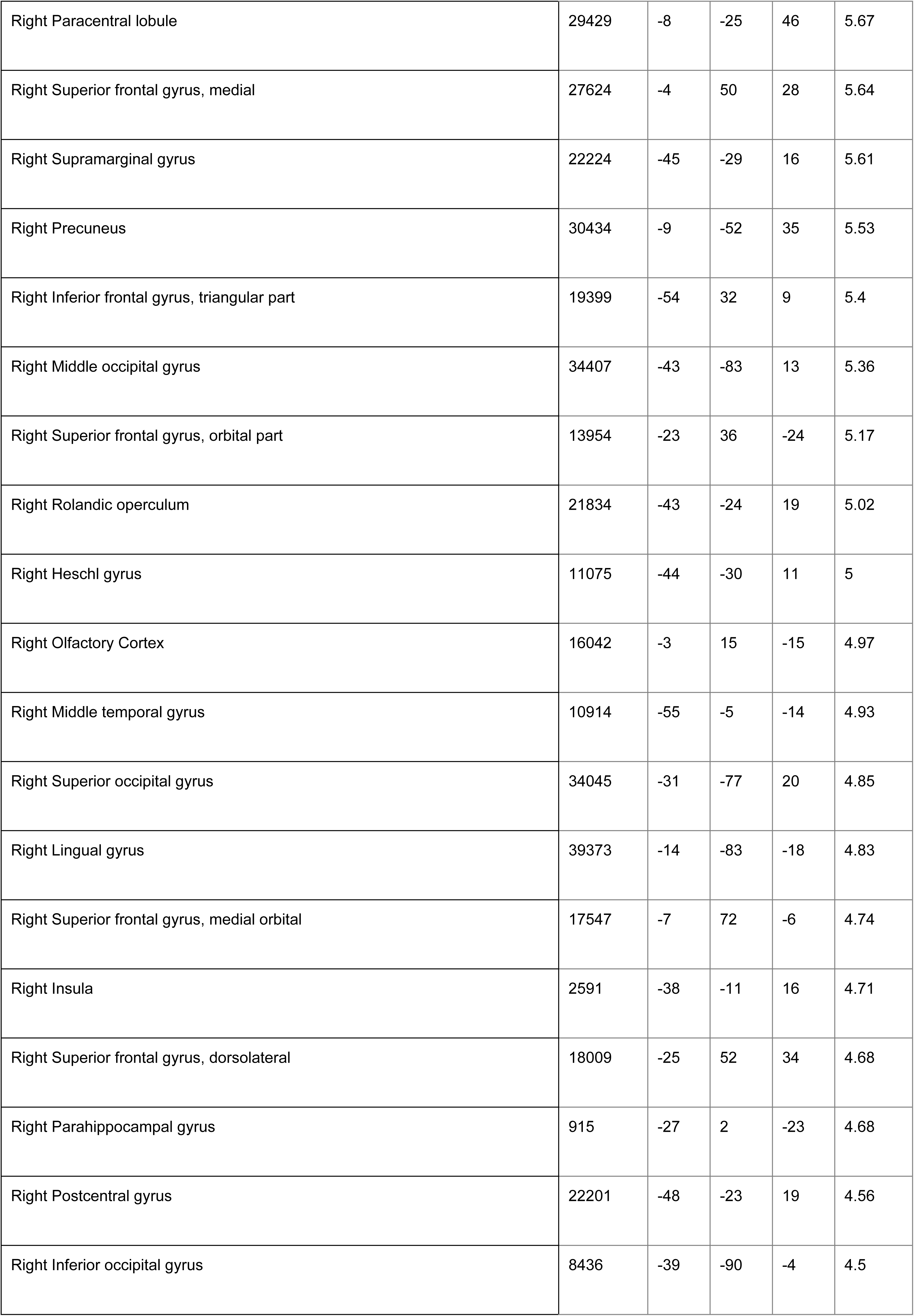
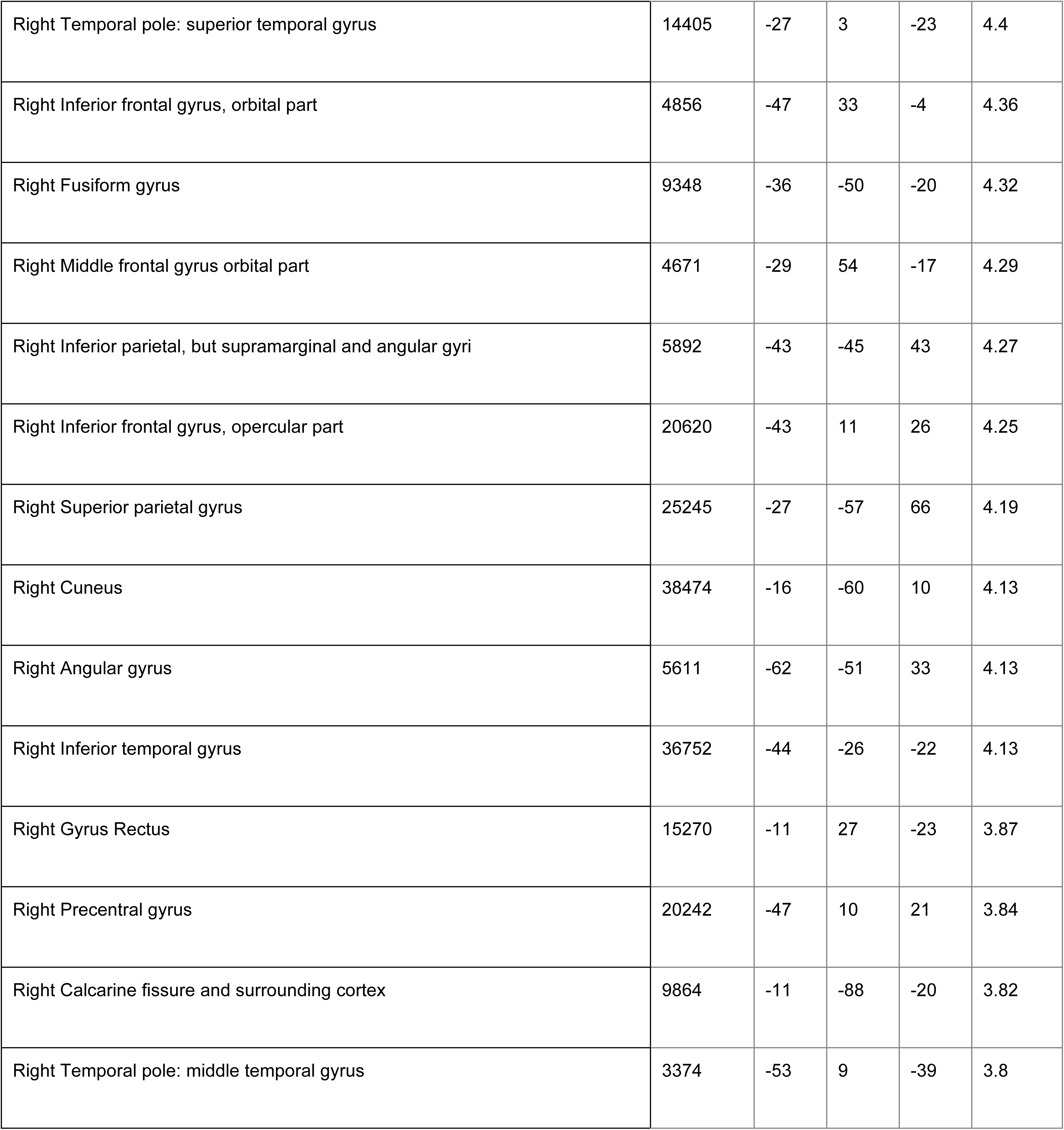
Left Striatum volume significant peaks of correlation with cortical thickness in HC.

**Supplementary Table 8.**
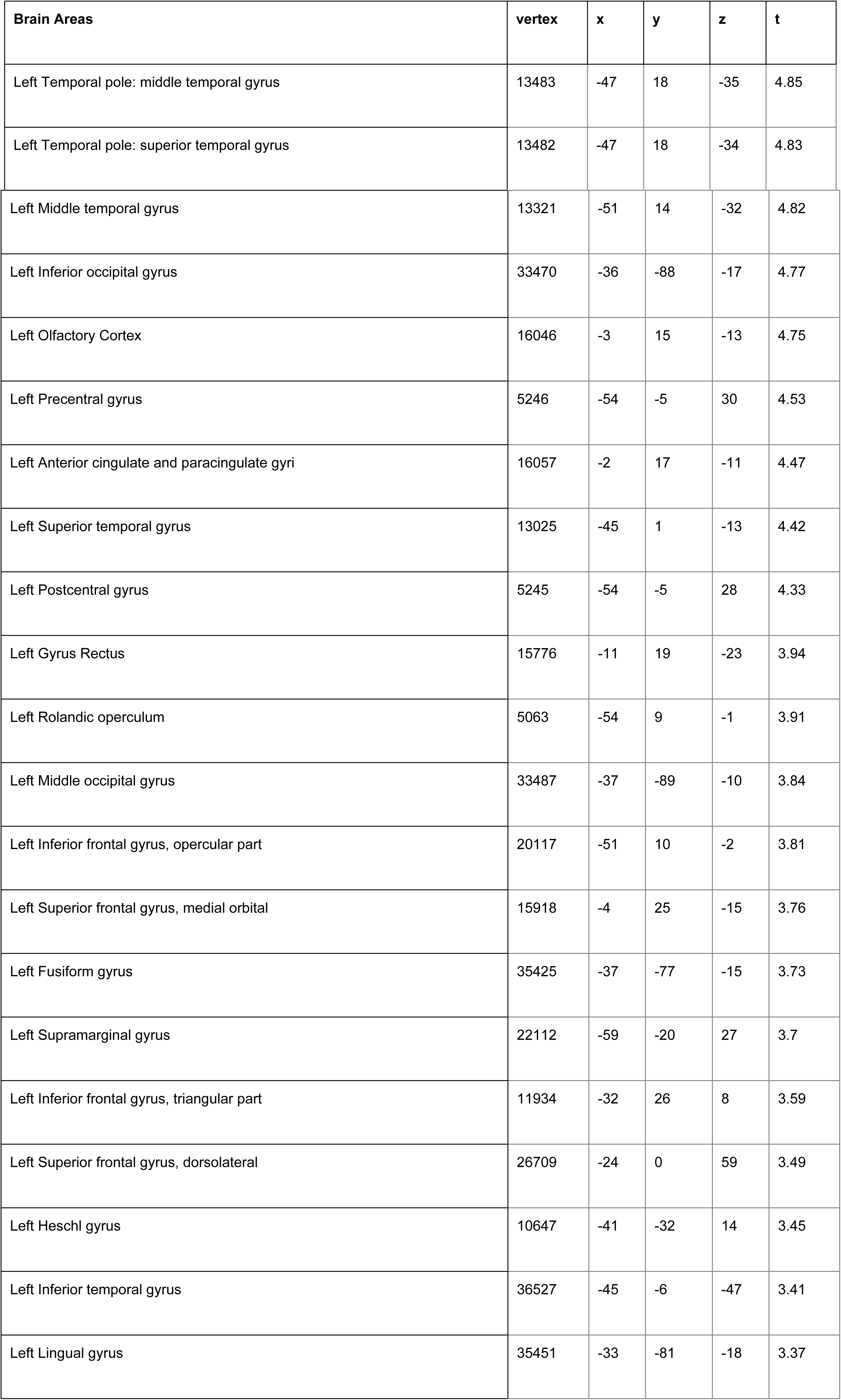
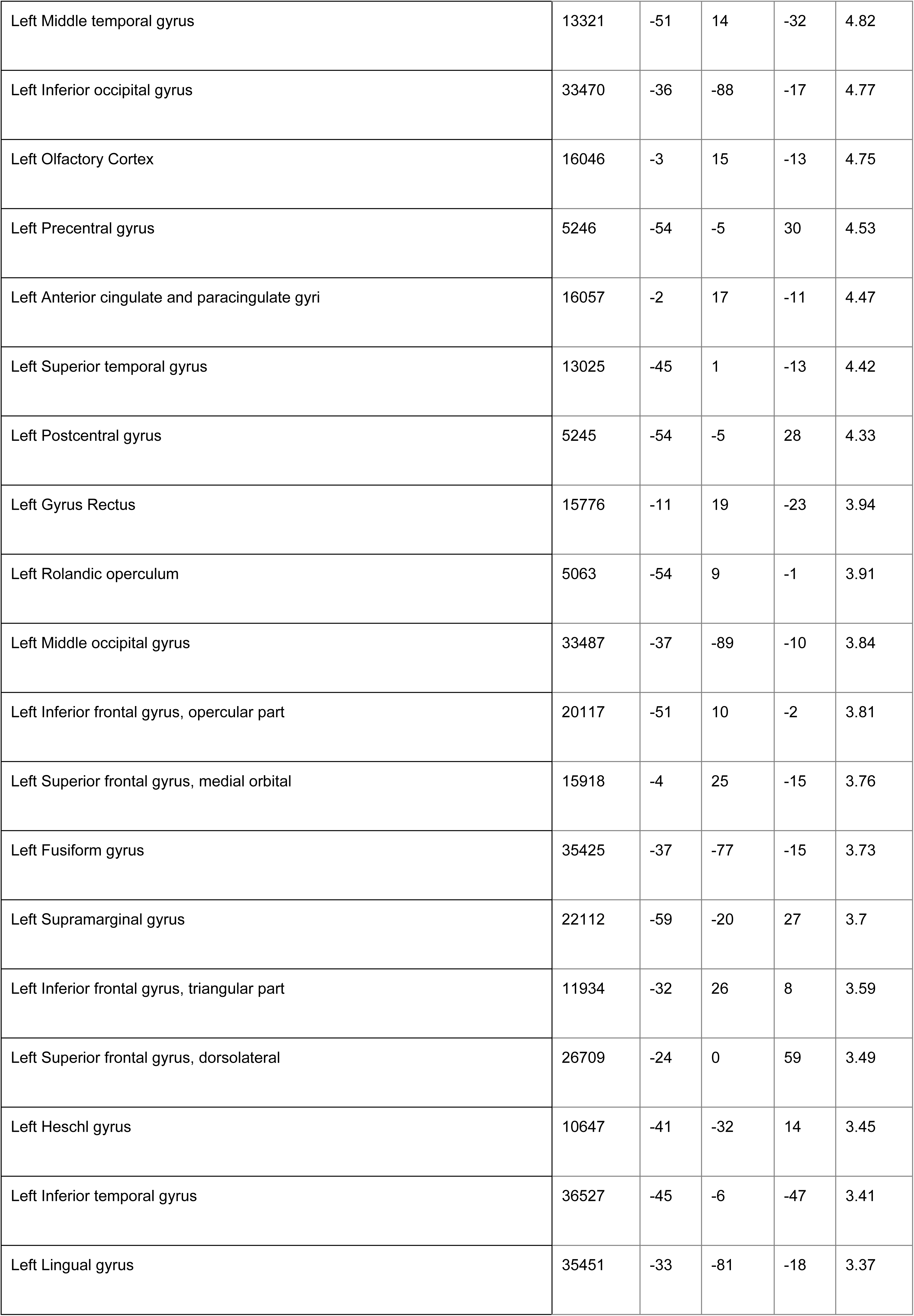
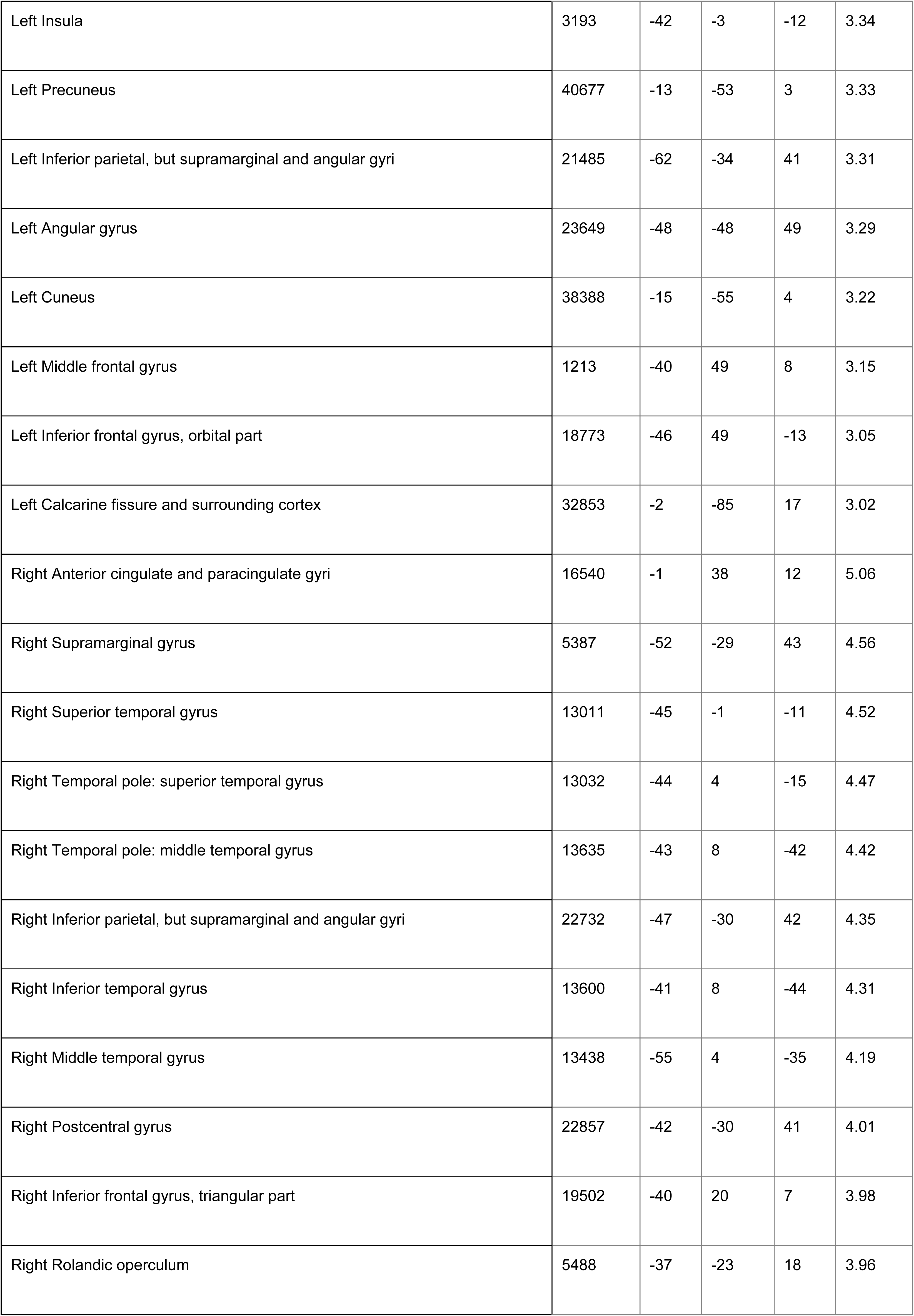
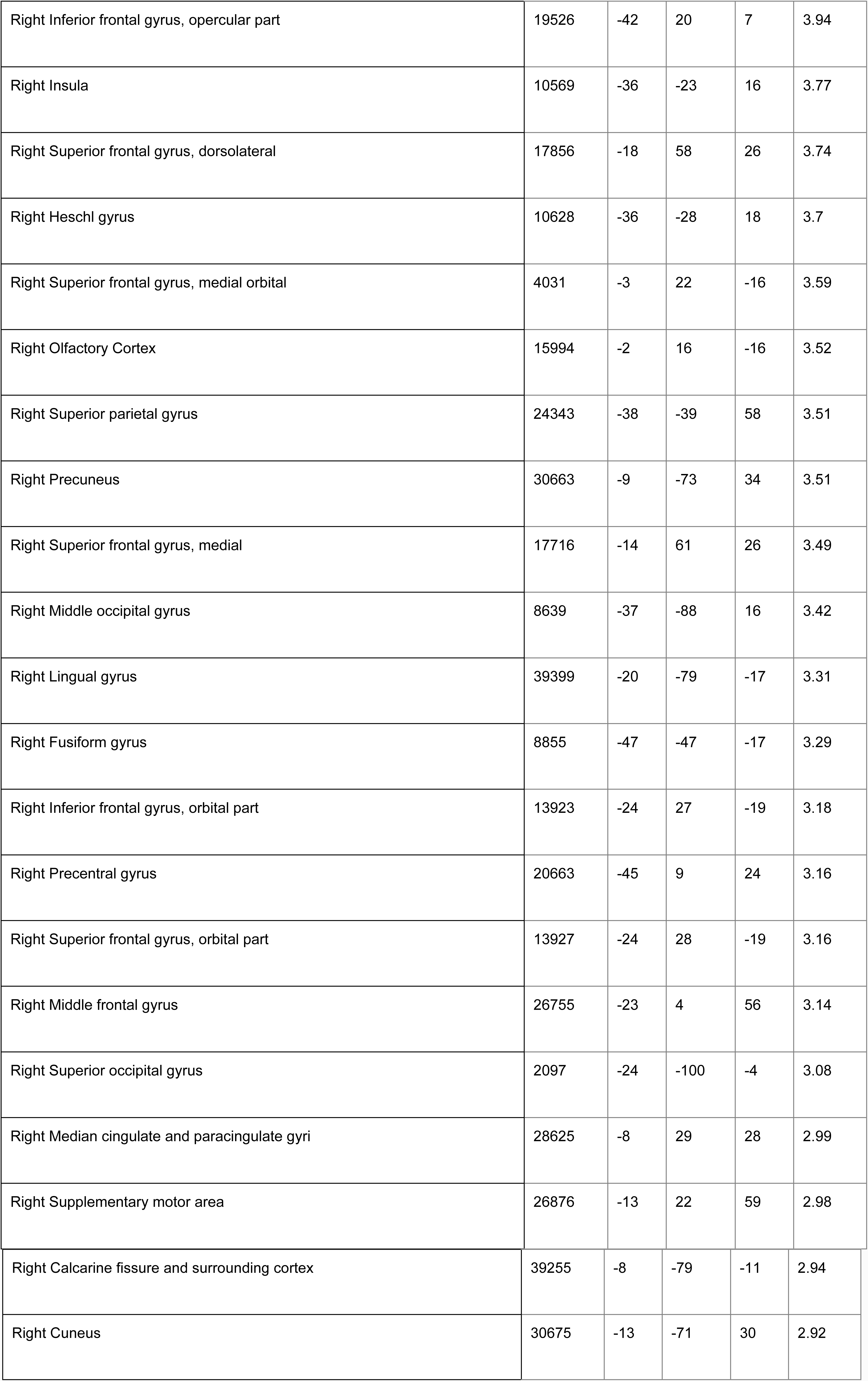
Left Striatum volume significant peaks of correlation with cortical thickness in AD.

**Supplementary Table 9.** Right Striatum volume significant peaks of correlation with cortical thickness in HC.

| Brain Areas | vertex | x | y | z | t |
| --- | --- | --- | --- | --- | --- |
| Left Calcarine fissure and surrounding cortex | 33295 | -11 | -92 | -19 | 6.02 |
| Left Precuneus | 30561 | -1 | -64 | 30 | 5.86 |
| Left Superior frontal gyrus, medial orbital | 16899 | -5 | 56 | -14 | 5.85 |
| Left Lingual gyrus | 39387 | -12 | -87 | -20 | 5.83 |
| Left Median cingulate and paracingulate gyri | 30717 | -5 | -41 | 48 | 5.53 |
| Left Inferior frontal gyrus, triangular part | 19949 | -43 | 30 | 21 | 5.45 |
| Left Inferior frontal gyrus, opercular part | 20044 | -48 | 13 | 24 | 5.33 |
| Left Middle frontal gyrus | 19179 | -40 | 32 | 19 | 5.29 |
| Left Gyrus Rectus | 4261 | -4 | 56 | -15 | 5.2 |
| Left Middle occipital gyrus | 8596 | -46 | -70 | 16 | 5.15 |
| Left Fusiform gyrus | 9424 | -37 | -57 | -20 | 4.67 |
| Left Inferior occipital gyrus | 35622 | -45 | -66 | -5 | 4.66 |
| Left Supplementary motor area | 7023 | -6 | 5 | 46 | 4.63 |
| Left Posterior cingulate gyrus | 30229 | -3 | -53 | 19 | 4.61 |
| Left Precentral gyrus | 20657 | -44 | 9 | 24 | 4.49 |
| Left Superior frontal gyrus, dorsolateral | 26093 | -21 | 27 | 59 | 4.28 |
| Left Superior occipital gyrus | 32420 | -29 | -86 | 29 | 4.28 |
| Left Superior frontal gyrus, medial | 444 | -8 | 53 | 43 | 4.27 |
| Left Middle temporal gyrus | 34872 | -51 | -54 | 15 | 4.21 |
| Left Superior parietal gyrus | 25145 | -31 | -55 | 65 | 4.2 |
| Left Insula | 10525 | -39 | -17 | 5 | 4.17 |
| Left Cuneus | 9614 | -18 | -53 | 1 | 4.16 |
| Left Postcentral gyrus | 25000 | -27 | -37 | 62 | 4.1 |
| Left Inferior temporal gyrus | 35914 | -61 | -56 | -8 | 4.1 |
| Left Rolandic operculum | 21690 | -39 | -12 | 17 | 3.98 |
| Left Heschl gyrus | 2642 | -39 | -19 | 4 | 3.96 |
| Left Angular gyrus | 5941 | -45 | -48 | 47 | 3.91 |
| Left Superior temporal gyrus | 2616 | -40 | -18 | 0 | 3.9 |
| Left Inferior parietal, but supramarginal and angular gyri | 23644 | -45 | -47 | 46 | 3.87 |
| Left Middle frontal gyrus orbital part | 4774 | -37 | 58 | 3 | 3.85 |
| Left Olfactory Cortex | 16045 | -3 | 16 | -14 | 3.81 |
| Left Supramarginal gyrus | 22367 | -51 | -38 | 24 | 3.81 |
| Left Anterior cingulate and paracingulate gyri | 15430 | -5 | 25 | -14 | 3.67 |
| Left Paracentral lobule | 30989 | -13 | -44 | 64 | 3.27 |
| Left Superior frontal gyrus, orbital part | 15200 | -11 | 37 | -23 | 2.99 |
| Left Temporal pole: superior temporal gyrus | 3307 | -44 | 18 | -19 | 2.73 |
| Left Inferior frontal gyrus, orbital part | 4808 | -42 | 46 | -2 | 2.68 |
| Left Parahippocampal gyrus | 3614 | -27 | 0 | -23 | 2.61 |
| Left Temporal pole: middle temporal gyrus | 3340 | -48 | 12 | -30 | 2.47 |
| Right Median cingulate and paracingulate gyri | 29254 | -11 | -15 | 44 | 6.22 |
| Right Supplementary motor area | 27989 | -8 | 2 | 46 | 5.85 |
| Right Superior temporal gyrus | 10884 | -59 | -8 | -7 | 5.74 |
| Right Posterior cingulate gyrus | 7574 | -4 | -48 | 35 | 5.73 |
| Right Middle frontal gyrus | 1166 | -33 | 44 | 28 | 5.65 |
| Right Middle occipital gyrus | 34407 | -43 | -83 | 13 | 5.56 |
| Right Inferior frontal gyrus, triangular part | 19399 | -54 | 32 | 9 | 5.55 |
| Right Paracentral lobule | 29429 | -8 | -25 | 46 | 5.49 |
| Right Anterior cingulate and paracingulate gyri | 28019 | -4 | 48 | 27 | 5.35 |
| Right Precuneus | 40664 | -6 | -48 | 2 | 5.34 |
| Right Superior frontal gyrus, medial | 27624 | -4 | 50 | 28 | 5.26 |
| Right Supramarginal gyrus | 22224 | -45 | -29 | 16 | 5.25 |
| Right Olfactory Cortex | 16042 | -3 | 15 | -15 | 5.16 |
| Right Superior frontal gyrus, orbital part | 3507 | -25 | 33 | -22 | 5.13 |
| Right Middle temporal gyrus | 10914 | -55 | -5 | -14 | 5.05 |
| Right Lingual gyrus | 39373 | -14 | -83 | -18 | 4.91 |
| Right Superior occipital gyrus | 34019 | -26 | -84 | 18 | 4.78 |
| Right Insula | 12731 | -36 | 23 | -2 | 4.77 |
| Right Rolandic operculum | 21834 | -43 | -24 | 19 | 4.73 |
| Right Heschl gyrus | 11075 | -44 | -30 | 11 | 4.58 |
| Right Superior frontal gyrus, medial orbital | 17547 | -7 | 72 | -6 | 4.55 |
| Right Superior frontal gyrus, dorsolateral | 27271 | -25 | 51 | 37 | 4.44 |
| Right Inferior occipital gyrus | 8436 | -39 | -90 | -4 | 4.44 |
| Right Fusiform gyrus | 9422 | -35 | -54 | -19 | 4.41 |
| Right Parahippocampal gyrus | 915 | -27 | 2 | -23 | 4.36 |
| Right Inferior frontal gyrus, opercular part | 5161 | -46 | 14 | 28 | 4.21 |
| Right Postcentral gyrus | 22201 | -48 | -23 | 19 | 4.18 |
| Right Inferior frontal gyrus, orbital part | 19170 | -50 | 35 | -7 | 4.16 |
| Right Inferior temporal gyrus | 36752 | -44 | -26 | -22 | 4.14 |
| Right Temporal pole: superior temporal gyrus | 14367 | -27 | 2 | -24 | 4.08 |
| Right Cuneus | 38474 | -16 | -60 | 10 | 3.98 |
| Right Middle frontal gyrus orbital part | 18538 | -23 | 51 | -14 | 3.94 |
| Right Inferior parietal, but supramarginal and angular gyri | 5892 | -43 | -45 | 43 | 3.89 |
| Right Calcarine fissure and surrounding cortex | 9864 | -11 | -88 | -20 | 3.82 |
| Right Angular gyrus | 5611 | -62 | -51 | 33 | 3.81 |
| Right Superior parietal gyrus | 25245 | -27 | -57 | 66 | 3.79 |
| Right Precentral gyrus | 20433 | -40 | 4 | 38 | 3.65 |
| Right Gyrus Rectus | 15270 | -11 | 27 | -23 | 3.46 |
| Right Temporal pole: middle temporal gyrus | 3374 | -53 | 9 | -39 | 3.45 |

**Supplementary Table 10.** Right Striatum volume significant peaks of correlation with cortical thickness in AD.

| Brain Areas | vertex | x | y | z | t |
| --- | --- | --- | --- | --- | --- |
| Left Olfactory Cortex | 16045 | -3 | 16 | -14 | 4.72 |
| Left Temporal pole: superior temporal gyrus | 13190 | -48 | 18 | -33 | 4.69 |
| Left Temporal pole: middle temporal gyrus | 13483 | -47 | 18 | -35 | 4.63 |
| Left Middle temporal gyrus | 13321 | -51 | 14 | -32 | 4.58 |
| Left Superior temporal gyrus | 3199 | -44 | 0 | -14 | 4.57 |
| Left Inferior occipital gyrus | 33470 | -36 | -88 | -17 | 4.42 |
| Left Anterior cingulate and paracingulate gyri | 16057 | -2 | 17 | -11 | 4.25 |
| Left Precentral gyrus | 20860 | -55 | -4 | 30 | 4.2 |
| Left Gyrus Rectus | 15776 | -11 | 19 | -23 | 4.08 |
| Left Postcentral gyrus | 5245 | -54 | -5 | 28 | 4.03 |
| Left Superior frontal gyrus, medial orbital | 1018 | -3 | 24 | -16 | 3.81 |
| Left Inferior frontal gyrus, triangular part | 11946 | -31 | 26 | 8 | 3.69 |
| Left Supramarginal gyrus | 21935 | -62 | -35 | 34 | 3.66 |
| Left Fusiform gyrus | 35425 | -37 | -77 | -15 | 3.52 |
| Left Insula | 3193 | -42 | -3 | -12 | 3.51 |
| Left Middle occipital gyrus | 33487 | -37 | -89 | -10 | 3.48 |
| Left Rolandic operculum | 5063 | -54 | 9 | -1 | 3.45 |
| Left Inferior frontal gyrus, opercular part | 20117 | -51 | 10 | -2 | 3.35 |
| Left Inferior parietal, but supramarginal and angular gyri | 21485 | -62 | -34 | 41 | 3.33 |
| Right Anterior cingulate and paracingulate gyri | 16540 | -1 | 38 | 12 | 5.24 |
| Right Supramarginal gyrus | 5387 | -52 | -29 | 43 | 4.55 |
| Right Superior temporal gyrus | 12707 | -44 | 1 | -14 | 4.45 |
| Right Temporal pole: superior temporal gyrus | 13034 | -44 | 4 | -16 | 4.38 |
| Right Inferior parietal, but supramarginal and angular gyri | 22732 | -47 | -30 | 42 | 4.3 |
| Right Temporal pole: middle temporal gyrus | 3429 | -42 | 9 | -41 | 4.28 |
| Right Inferior temporal gyrus | 13600 | -41 | 8 | -44 | 4.19 |
| Right Middle temporal gyrus | 13438 | -55 | 4 | -35 | 4.14 |
| Right Inferior frontal gyrus, triangular part | 19502 | -40 | 20 | 7 | 3.96 |
| Right Postcentral gyrus | 21313 | -51 | -26 | 38 | 3.96 |
| Right Rolandic operculum | 21847 | -39 | -21 | 17 | 3.93 |
| Right Inferior frontal gyrus, opercular part | 19526 | -42 | 20 | 7 | 3.89 |
| Right Superior frontal gyrus, dorsolateral | 17856 | -18 | 58 | 26 | 3.78 |
| Right Heschl gyrus | 10628 | -36 | -28 | 18 | 3.68 |
| Right Olfactory Cortex | 15994 | -2 | 16 | -16 | 3.64 |
| Right Insula | 10569 | -36 | -23 | 16 | 3.61 |
| Right Middle occipital gyrus | 8639 | -37 | -88 | 16 | 3.59 |
| Right Superior parietal gyrus | 24343 | -38 | -39 | 58 | 3.54 |
| Right Superior frontal gyrus, medial orbital | 4031 | -3 | 22 | -16 | 3.48 |
| Right Precuneus | 7939 | -11 | -76 | 49 | 3.46 |
| Right Superior frontal gyrus, medial | 4460 | -16 | 59 | 26 | 3.42 |
| Right Precentral gyrus | 20661 | -46 | 9 | 25 | 3.38 |
| Right Inferior frontal gyrus, orbital part | 13923 | -24 | 27 | -19 | 3.26 |
| Right Lingual gyrus | 39399 | -20 | -79 | -17 | 3.24 |
| Right Superior frontal gyrus, orbital part | 13927 | -24 | 28 | -19 | 3.23 |
| Right Fusiform gyrus | 13717 | -28 | 2 | -43 | 3.21 |
| Right Middle frontal gyrus | 26755 | -23 | 4 | 56 | 3.15 |

**Supplementary Table 11.** Left nucleus accumbens volume significant peaks of correlation with cortical thickness for ALL participants.

| Brain Areas | vertex | x | y | z | t |
| --- | --- | --- | --- | --- | --- |
| Left Calcarine fissure and surrounding cortex | 33204 | -10 | -92 | -19 | 5.92 |
| Left Lingual gyrus | 39386 | -10 | -86 | -19 | 5.25 |
| Left Olfactory Cortex | 16046 | -3 | 15 | -13 | 5.09 |
| Left Anterior cingulate and paracingulate gyri | 16057 | -2 | 17 | -11 | 4.84 |
| Left Cuneus | 38621 | -15 | -64 | 7 | 4.64 |
| Left Gyrus Rectus | 16944 | -2 | 60 | -18 | 4.6 |
| Left Superior frontal gyrus, medial orbital | 16951 | -2 | 60 | -14 | 4.39 |
| Left Superior parietal gyrus | 6284 | -27 | -37 | 61 | 4.17 |
| Left Superior temporal gyrus | 22384 | -45 | -34 | 17 | 4.15 |
| Left Postcentral gyrus | 24999 | -28 | -37 | 62 | 4.12 |
| Left Middle frontal gyrus | 25975 | -23 | 26 | 41 | 4.08 |
| Left Superior frontal gyrus, dorsolateral | 25993 | -22 | 27 | 41 | 3.96 |
| Left Inferior frontal gyrus, opercular part | 4909 | -38 | 17 | 5 | 3.95 |
| Left Middle temporal gyrus | 13324 | -52 | 10 | -28 | 3.89 |
| Left Rolandic operculum | 328 | -48 | 5 | 1 | 3.88 |
| Left Inferior frontal gyrus, triangular part | 19219 | -50 | 33 | 13 | 3.87 |
| Left Heschl gyrus | 11076 | -45 | -29 | 11 | 3.82 |
| Left Precuneus | 38525 | -17 | -60 | 15 | 3.8 |
| Left Superior frontal gyrus, medial | 28222 | -6 | 32 | 38 | 3.79 |
| Left Middle occipital gyrus | 33446 | -33 | -93 | -10 | 3.77 |
| Left Temporal pole: superior temporal gyrus | 3342 | -53 | 7 | -23 | 3.75 |
| Left Inferior occipital gyrus | 33441 | -32 | -94 | -12 | 3.74 |
| Left Supramarginal gyrus | 5586 | -45 | -34 | 18 | 3.73 |
| Left Parahippocampal gyrus | 38021 | -29 | -35 | -6 | 3.66 |
| Left Supplementary motor area | 28296 | -7 | 21 | 47 | 3.64 |
| Left Superior frontal gyrus, orbital part | 17132 | -8 | 65 | -20 | 3.63 |
| Left Inferior temporal gyrus | 35934 | -60 | -50 | -13 | 3.6 |
| Left Angular gyrus | 23298 | -48 | -54 | 27 | 3.52 |
| Left Insula | 12064 | -32 | 18 | 11 | 3.51 |
| Left Temporal pole: middle temporal gyrus | 3340 | -48 | 12 | -30 | 3.47 |
| Left Precentral gyrus | 26763 | -23 | -4 | 54 | 3.45 |
| Left Median cingulate and paracingulate gyri | 7483 | -5 | -18 | 44 | 3.38 |
| Left Inferior parietal, but supramarginal and angular gyri | 6032 | -27 | -65 | 35 | 3.07 |
| Left Fusiform gyrus | 9175 | -42 | -15 | -26 | 3.05 |
| Left Middle frontal gyrus orbital part | 18981 | -33 | 58 | 3 | 2.99 |
| Left Superior occipital gyrus | 24141 | -27 | -66 | 32 | 2.95 |
| Left Inferior frontal gyrus, orbital part | 18643 | -39 | 52 | -13 | 2.94 |
| Left Paracentral lobule | 30994 | -12 | -44 | 65 | 2.56 |
| Left Posterior cingulate gyrus | 40909 | -2 | -52 | 16 | 2.48 |
| Right Superior temporal gyrus | 11384 | -65 | -15 | -3 | 5.75 |
| Right Median cingulate and paracingulate gyri | 29746 | -11 | -20 | 41 | 5.45 |
| Right Middle temporal gyrus | 9116 | -52 | -46 | 9 | 5.4 |
| Right Temporal pole: middle temporal gyrus | 853 | -52 | 13 | -40 | 5.38 |
| Right Lingual gyrus | 39369 | -15 | -82 | -18 | 5.27 |
| Right Supramarginal gyrus | 22416 | -54 | -37 | 20 | 5.08 |
| Right Middle frontal gyrus | 19591 | -31 | 38 | 30 | 5.04 |
| Right Inferior frontal gyrus, triangular part | 19366 | -38 | 26 | 5 | 5.02 |
| Right Heschl gyrus | 11311 | -57 | -14 | 7 | 4.91 |
| Right Olfactory Cortex | 15923 | -7 | 7 | -18 | 4.88 |
| Right Inferior temporal gyrus | 13591 | -44 | 6 | -42 | 4.84 |
| Right Precuneus | 31620 | -10 | -76 | 53 | 4.73 |
| Right Insula | 10537 | -39 | -19 | 5 | 4.66 |
| Right Superior parietal gyrus | 31652 | -12 | -73 | 51 | 4.64 |
| Right Superior frontal gyrus, dorsolateral | 17921 | -23 | 58 | 19 | 4.57 |
| Right Anterior cingulate and paracingulate gyri | 16045 | -3 | 16 | -14 | 4.57 |
| Right Angular gyrus | 24041 | -28 | -67 | 33 | 4.52 |
| Right Inferior parietal, but supramarginal and angular gyri | 390 | -27 | -65 | 33 | 4.45 |
| Right Calcarine fissure and surrounding cortex | 39283 | -9 | -83 | -15 | 4.42 |
| Right Cuneus | 31925 | -16 | -65 | 19 | 4.41 |
| Right Middle occipital gyrus | 24060 | -28 | -67 | 32 | 4.41 |
| Right Postcentral gyrus | 6134 | -41 | -34 | 58 | 4.31 |
| Right Posterior cingulate gyrus | 29964 | 0 | -33 | 33 | 4.17 |
| Right Temporal pole: superior temporal gyrus | 13495 | -46 | 21 | -35 | 4.1 |
| Right Superior occipital gyrus | 34019 | -26 | -84 | 18 | 4.02 |
| Right Inferior frontal gyrus, opercular part | 4908 | -41 | 20 | 7 | 3.95 |
| Right Superior frontal gyrus, medial | 4440 | -4 | 52 | 25 | 3.94 |
| Right Fusiform gyrus | 37897 | -25 | -76 | -13 | 3.85 |
| Right Supplementary motor area | 7386 | -7 | -21 | 45 | 3.81 |
| Right Paracentral lobule | 29424 | -8 | -24 | 46 | 3.79 |
| Right Precentral gyrus | 21025 | -39 | -18 | 43 | 3.77 |
| Right Rolandic operculum | 5489 | -41 | -24 | 19 | 3.76 |
| Right Inferior frontal gyrus, orbital part | 19169 | -51 | 38 | -7 | 3.62 |
| Right Gyrus Rectus | 15273 | -12 | 25 | -22 | 3.56 |
| Right Superior frontal gyrus, orbital part | 15242 | -12 | 26 | -24 | 3.49 |
| Right Superior frontal gyrus, medial orbital | 76 | -6 | 70 | -11 | 3.35 |
| Right Inferior occipital gyrus | 33607 | -35 | -91 | -7 | 3.2 |
| Right Middle frontal gyrus orbital part | 18445 | -28 | 55 | -1 | 3.11 |
| Right Parahippocampal gyrus | 915 | -27 | 2 | -23 | 2.79 |

**Supplementary Table 12.**
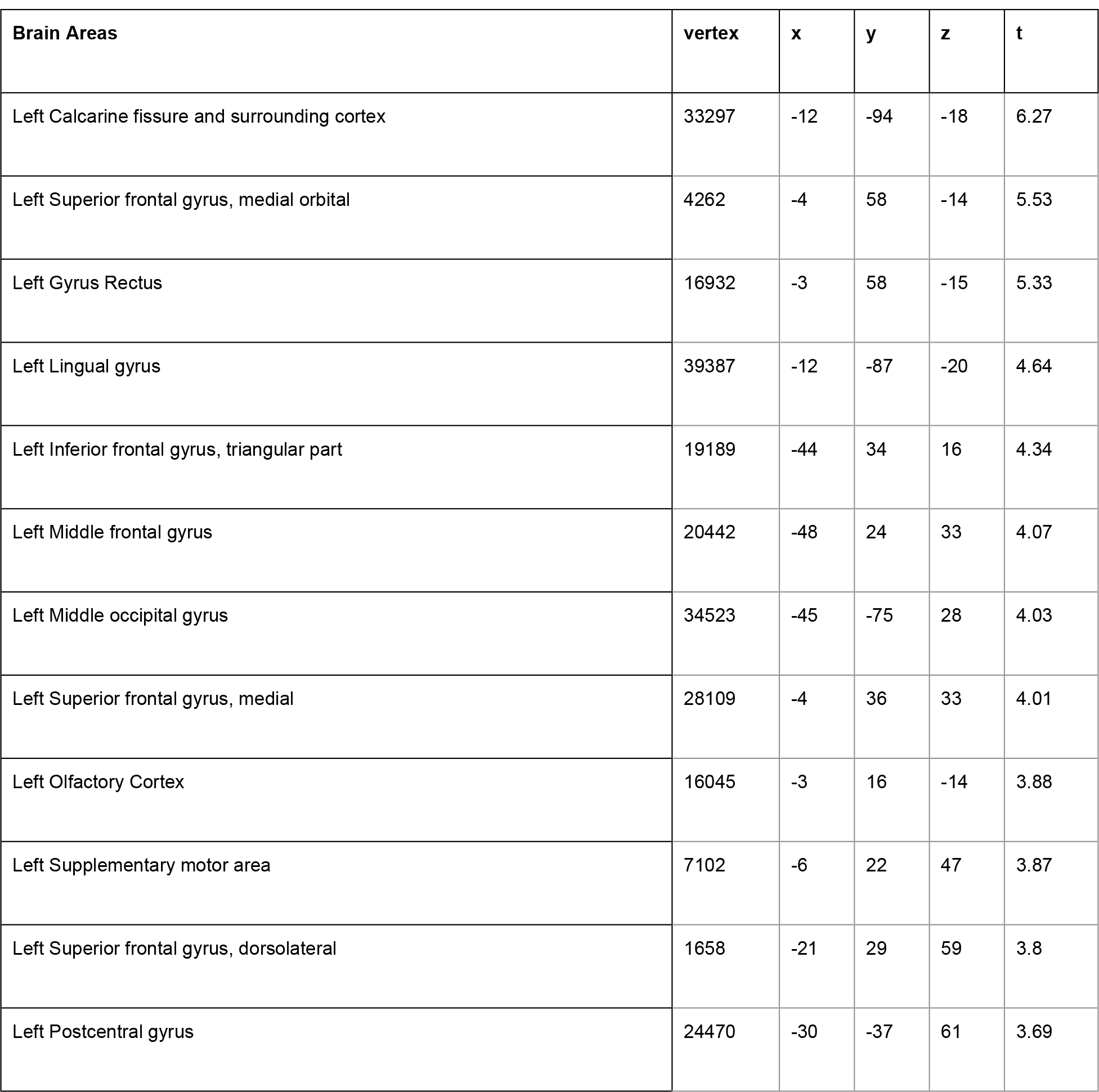
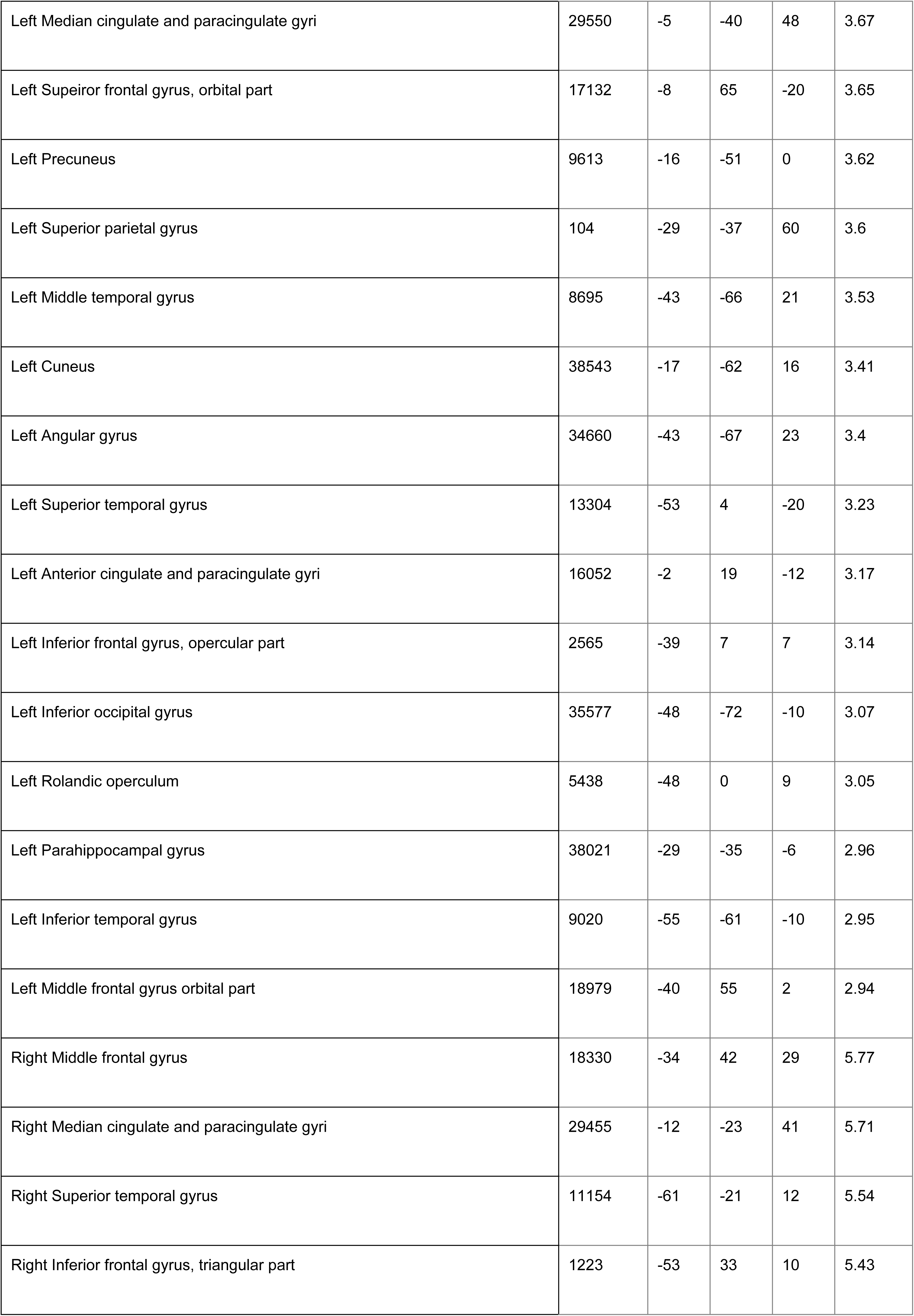
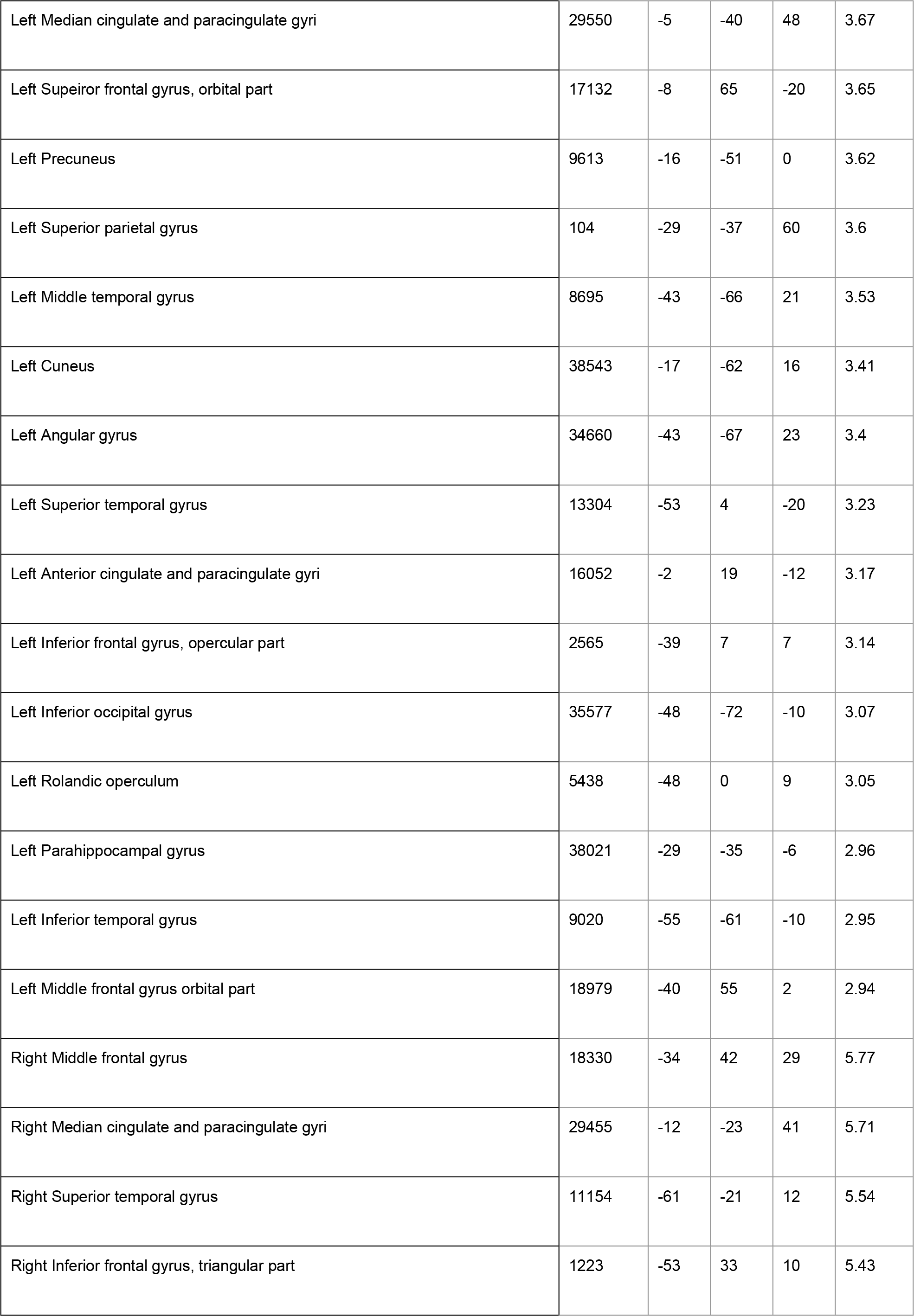
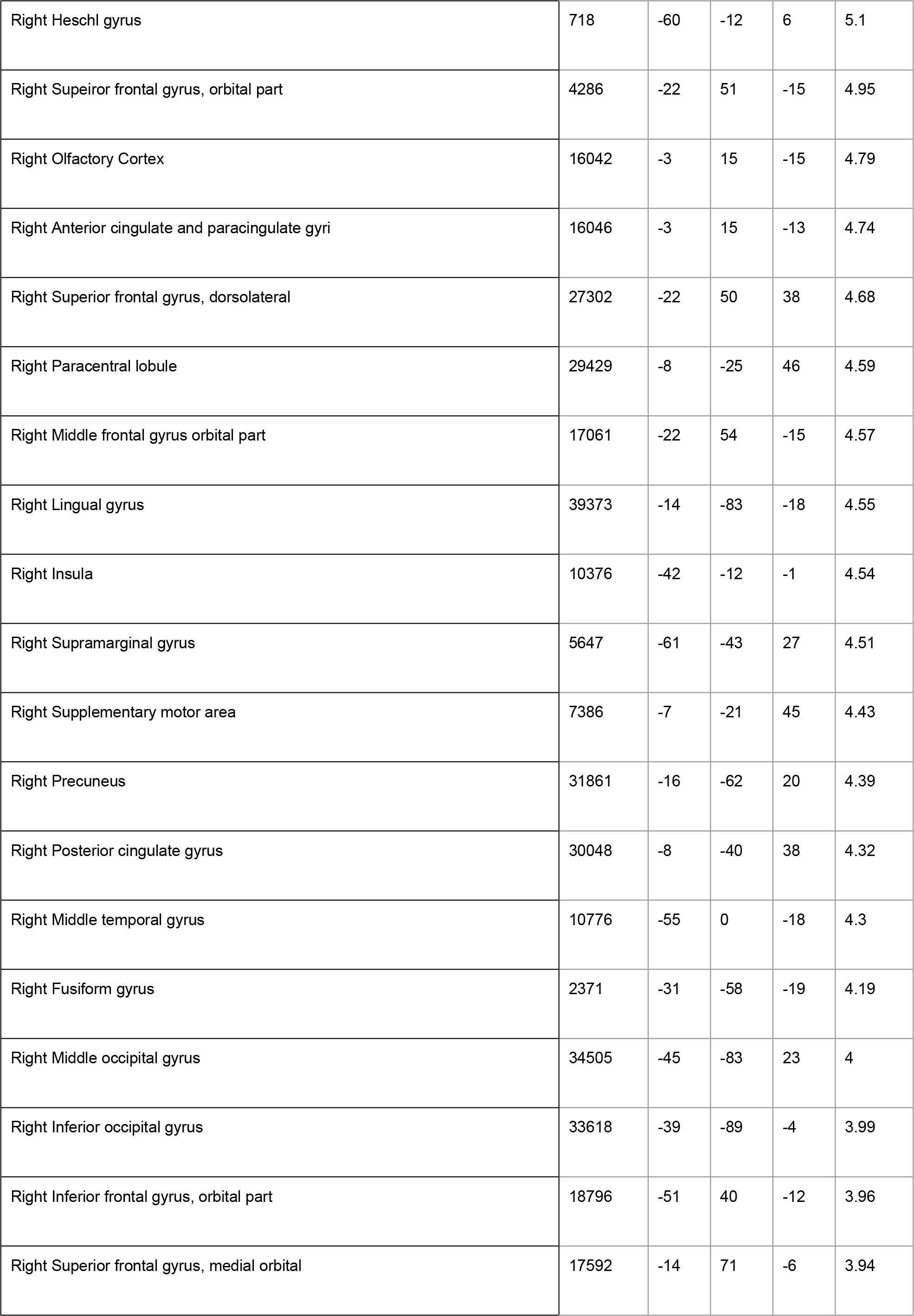
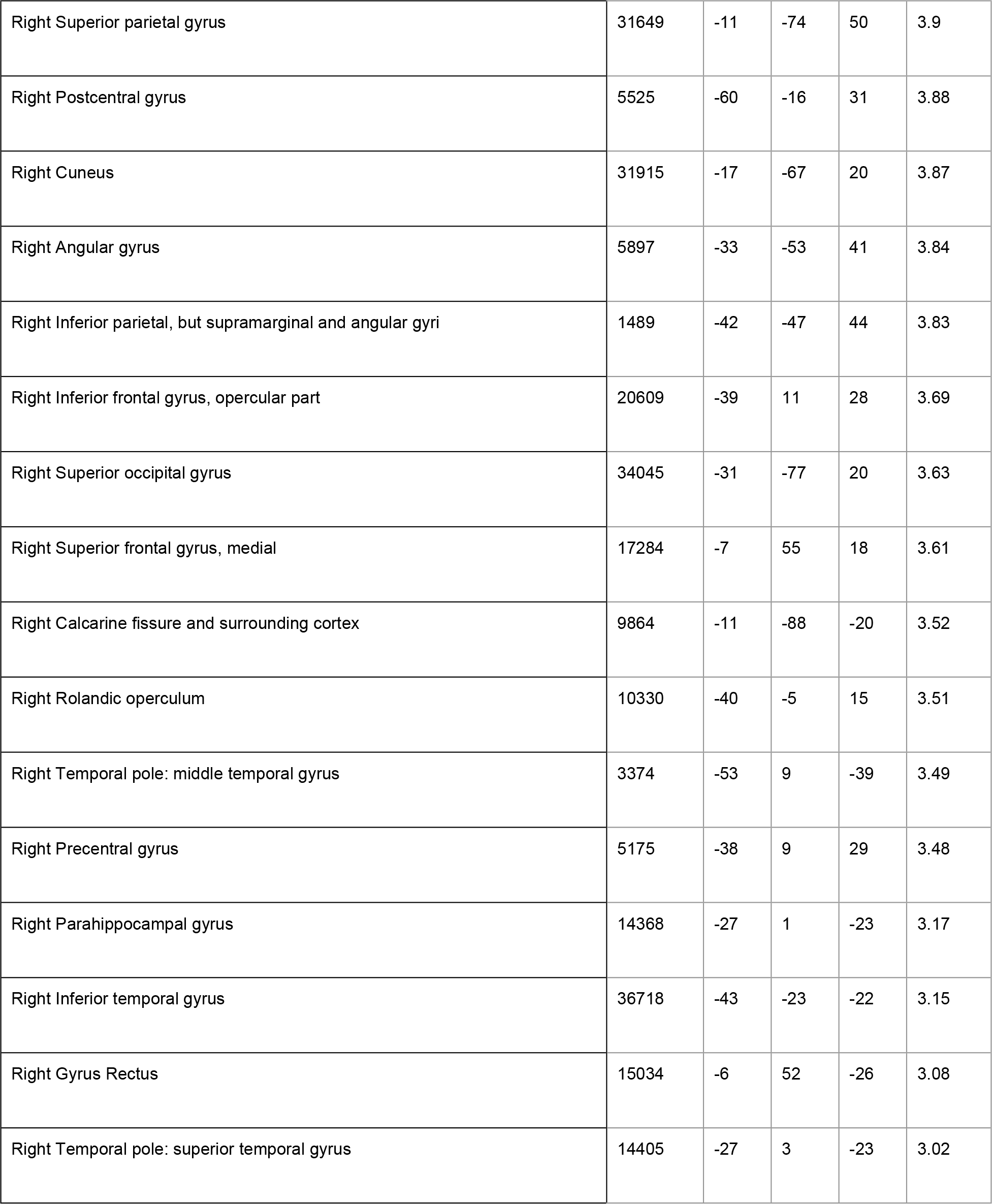
Left nucleus accumbens volume significant peaks of correlation with cortical thickness for HC.

| Brain Areas | vertex | x | y | z | t |
| --- | --- | --- | --- | --- | --- |
| Left Calcarine fissure and surrounding cortex | 33297 | -12 | -94 | -18 | 6.27 |
| Left Superior frontal gyrus, medial orbital | 4262 | -4 | 58 | -14 | 5.53 |
| Left Gyrus Rectus | 16932 | -3 | 58 | -15 | 5.33 |
| Left Lingual gyrus | 39387 | -12 | -87 | -20 | 4.64 |
| Left Inferior frontal gyrus, triangular part | 19189 | -44 | 34 | 16 | 4.34 |
| Left Middle frontal gyrus | 20442 | -48 | 24 | 33 | 4.07 |
| Left Middle occipital gyrus | 34523 | -45 | -75 | 28 | 4.03 |
| Left Superior frontal gyrus, medial | 28109 | -4 | 36 | 33 | 4.01 |
| Left Olfactory Cortex | 16045 | -3 | 16 | -14 | 3.88 |
| Left Supplementary motor area | 7102 | -6 | 22 | 47 | 3.87 |
| Left Superior frontal gyrus, dorsolateral | 1658 | -21 | 29 | 59 | 3.8 |
| Left Postcentral gyrus | 24470 | -30 | -37 | 61 | 3.69 |
| Left Median cingulate and paracingulate gyri | 29550 | -5 | -40 | 48 | 3.67 |
| Left Superior frontal gyrus, orbital part | 17132 | -8 | 65 | -20 | 3.65 |
| Left Precuneus | 9613 | -16 | -51 | 0 | 3.62 |
| Left Superior parietal gyrus | 104 | -29 | -37 | 60 | 3.6 |
| Left Middle temporal gyrus | 8695 | -43 | -66 | 21 | 3.53 |
| Left Cuneus | 38543 | -17 | -62 | 16 | 3.41 |
| Left Angular gyrus | 34660 | -43 | -67 | 23 | 3.4 |
| Left Superior temporal gyrus | 13304 | -53 | 4 | -20 | 3.23 |
| Left Anterior cingulate and paracingulate gyri | 16052 | -2 | 19 | -12 | 3.17 |
| Left Inferior frontal gyrus, opercular part | 2565 | -39 | 7 | 7 | 3.14 |
| Left Inferior occipital gyrus | 35577 | -48 | -72 | -10 | 3.07 |
| Left Rolandic operculum | 5438 | -48 | 0 | 9 | 3.05 |
| Left Parahippocampal gyrus | 38021 | -29 | -35 | -6 | 2.96 |
| Left Inferior temporal gyrus | 9020 | -55 | -61 | -10 | 2.95 |
| Left Middle frontal gyrus orbital part | 18979 | -40 | 55 | 2 | 2.94 |
| Right Middle frontal gyrus | 18330 | -34 | 42 | 29 | 5.77 |
| Right Median cingulate and paracingulate gyri | 29455 | -12 | -23 | 41 | 5.71 |
| Right Superior temporal gyrus | 11154 | -61 | -21 | 12 | 5.54 |
| Right Inferior frontal gyrus, triangular part | 1223 | -53 | 33 | 10 | 5.43 |
| Right Heschl gyrus | 718 | -60 | -12 | 6 | 5.1 |
| Right Superior frontal gyrus, orbital part | 4286 | -22 | 51 | -15 | 4.95 |
| Right Olfactory Cortex | 16042 | -3 | 15 | -15 | 4.79 |
| Right Anterior cingulate and paracingulate gyri | 16046 | -3 | 15 | -13 | 4.74 |
| Right Superior frontal gyrus, dorsolateral | 27302 | -22 | 50 | 38 | 4.68 |
| Right Paracentral lobule | 29429 | -8 | -25 | 46 | 4.59 |
| Right Middle frontal gyrus orbital part | 17061 | -22 | 54 | -15 | 4.57 |
| Right Lingual gyrus | 39373 | -14 | -83 | -18 | 4.55 |
| Right Insula | 10376 | -42 | -12 | -1 | 4.54 |
| Right Supramarginal gyrus | 5647 | -61 | -43 | 27 | 4.51 |
| Right Supplementary motor area | 7386 | -7 | -21 | 45 | 4.43 |
| Right Precuneus | 31861 | -16 | -62 | 20 | 4.39 |
| Right Posterior cingulate gyrus | 30048 | -8 | -40 | 38 | 4.32 |
| Right Middle temporal gyrus | 10776 | -55 | 0 | -18 | 4.3 |
| Right Fusiform gyrus | 2371 | -31 | -58 | -19 | 4.19 |
| Right Middle occipital gyrus | 34505 | -45 | -83 | 23 | 4 |
| Right Inferior occipital gyrus | 33618 | -39 | -89 | -4 | 3.99 |
| Right Inferior frontal gyrus, orbital part | 18796 | -51 | 40 | -12 | 3.96 |
| Right Superior frontal gyrus, medial orbital | 17592 | -14 | 71 | -6 | 3.94 |
| Right Superior parietal gyrus | 31649 | -11 | -74 | 50 | 3.9 |
| Right Postcentral gyrus | 5525 | -60 | -16 | 31 | 3.88 |
| Right Cuneus | 31915 | -17 | -67 | 20 | 3.87 |
| Right Angular gyrus | 5897 | -33 | -53 | 41 | 3.84 |
| Right Inferior parietal, but supramarginal and angular gyri | 1489 | -42 | -47 | 44 | 3.83 |
| Right Inferior frontal gyrus, opercular part | 20609 | -39 | 11 | 28 | 3.69 |
| Right Superior occipital gyrus | 34045 | -31 | -77 | 20 | 3.63 |
| Right Superior frontal gyrus, medial | 17284 | -7 | 55 | 18 | 3.61 |
| Right Calcarine fissure and surrounding cortex | 9864 | -11 | -88 | -20 | 3.52 |
| Right Rolandic operculum | 10330 | -40 | -5 | 15 | 3.51 |
| Right Temporal pole: middle temporal gyrus | 3374 | -53 | 9 | -39 | 3.49 |
| Right Precentral gyrus | 5175 | -38 | 9 | 29 | 3.48 |
| Right Parahippocampal gyrus | 14368 | -27 | 1 | -23 | 3.17 |
| Right Inferior temporal gyrus | 36718 | -43 | -23 | -22 | 3.15 |
| Right Gyrus Rectus | 15034 | -6 | 52 | -26 | 3.08 |
| Right Temporal pole: superior temporal gyrus | 14405 | -27 | 3 | -23 | 3.02 |

**Supplementary Table 13.** Right precommisural putamen volume significant peaks of correlation with cortical thickness for ALL participants.

| Brain Areas | vertex | x | y | z | t |
| --- | --- | --- | --- | --- | --- |
| Left Precuneus | 31341 | -3 | -69 | 45 | 4.94 |
| Left Middle frontal gyrus | 18965 | -32 | 56 | 7 | 4.74 |
| Left Lingual gyrus | 2475 | -13 | -86 | -19 | 4.43 |
| Left Insula | 2639 | -40 | -16 | 4 | 4.36 |
| Left Middle frontal gyrus orbital part | 18981 | -33 | 58 | 3 | 4.35 |
| Left Calcarine fissure and surrounding cortex | 39420 | -13 | -89 | -20 | 4.34 |
| Left Middle temporal gyrus | 13331 | -53 | 12 | -30 | 4.25 |
| Left Inferior occipital gyrus | 33458 | -32 | -94 | -14 | 4.21 |
| Left Temporal pole: superior temporal gyrus | 13188 | -49 | 16 | -30 | 4.2 |
| Left Temporal pole: middle temporal gyrus | 13285 | -48 | 13 | -30 | 4.09 |
| Left Inferior temporal gyrus | 35837 | -50 | -50 | -20 | 4.08 |
| Left Heschl gyrus | 2642 | -39 | -19 | 4 | 3.98 |
| Left Superior temporal gyrus | 22418 | -53 | -36 | 19 | 3.9 |
| Left Olfactory Cortex | 15928 | -5 | 9 | -18 | 3.88 |
| Left Precentral gyrus | 20887 | -53 | -5 | 30 | 3.78 |
| Left Superior frontal gyrus, dorsolateral | 6554 | -23 | 24 | 57 | 3.73 |
| Left Fusiform gyrus | 37217 | -46 | -42 | -21 | 3.73 |
| Left Inferior frontal gyrus, triangular part | 19050 | -40 | 45 | 7 | 3.7 |
| Left Median cingulate and paracingulate gyri | 29797 | -1 | -22 | 42 | 3.7 |
| Left Superior frontal gyrus, medial | 27542 | -7 | 55 | 42 | 3.61 |
| Left Superior frontal gyrus, medial orbital | 4262 | -4 | 58 | -14 | 3.61 |
| Left Gyrus Rectus | 15220 | -9 | 36 | -24 | 3.61 |
| Left Inferior frontal gyrus, opercular part | 20608 | -40 | 12 | 28 | 3.57 |
| Left Superior frontal gyrus, orbital part | 18442 | -27 | 54 | 1 | 3.55 |
| Left Middle occipital gyrus | 33446 | -33 | -93 | -10 | 3.53 |
| Left Supramarginal gyrus | 22355 | -53 | -39 | 25 | 3.51 |
| Left Postcentral gyrus | 20865 | -53 | -6 | 29 | 3.46 |
| Left Posterior cingulate gyrus | 40909 | -2 | -52 | 16 | 3.41 |
| Left Superior parietal gyrus | 31636 | -11 | -74 | 58 | 3.38 |
| Left Anterior cingulate and paracingulate gyri | 16057 | -2 | 17 | -11 | 3.37 |
| Left Cuneus | 32792 | -3 | -86 | 28 | 3.34 |
| Left Rolandic operculum | 21802 | -46 | -23 | 19 | 3.31 |
| Left Inferior parietal, but supramarginal and angular gyri | 23463 | -33 | -55 | 42 | 3.11 |
| Left Superior occipital gyrus | 33892 | -28 | -84 | 23 | 3.07 |
| Left Supplementary motor area | 6665 | -16 | 16 | 66 | 2.99 |
| Left Angular gyrus | 23487 | -35 | -57 | 43 | 2.82 |
| Left Inferior frontal gyrus, orbital part | 4808 | -42 | 46 | -2 | 2.6 |
| Right Supramarginal gyrus | 22409 | -50 | -36 | 20 | 4.78 |
| Right Median cingulate and paracingulate gyri | 7258 | -2 | 3 | 43 | 4.76 |
| Right Superior temporal gyrus | 22390 | -50 | -36 | 19 | 4.53 |
| Right Lingual gyrus | 624 | -18 | -80 | -19 | 4.51 |
| Right Inferior frontal gyrus, triangular part | 19491 | -41 | 22 | 7 | 4.43 |
| Right Anterior cingulate and paracingulate gyri | 16045 | -3 | 16 | -14 | 4.39 |
| Right Olfactory Cortex | 16042 | -3 | 15 | -15 | 4.26 |
| Right Precuneus | 30243 | -2 | -55 | 18 | 4.26 |
| Right Rolandic operculum | 21834 | -43 | -24 | 19 | 4.23 |
| Right Insula | 10531 | -38 | -19 | 6 | 4.19 |
| Right Supplementary motor area | 28014 | -9 | 2 | 44 | 4 |
| Right Inferior frontal gyrus, opercular part | 4904 | -38 | 19 | 7 | 3.99 |
| Right Heschl gyrus | 10543 | -38 | -20 | 5 | 3.96 |
| Right Posterior cingulate gyrus | 30231 | -5 | -53 | 22 | 3.86 |
| Right Temporal pole: middle temporal gyrus | 13419 | -54 | 10 | -37 | 3.85 |
| Right Middle temporal gyrus | 13409 | -56 | 9 | -36 | 3.82 |
| Right Middle occipital gyrus | 2155 | -35 | -88 | 22 | 3.74 |
| Right Inferior temporal gyrus | 35845 | -49 | -53 | -17 | 3.71 |
| Right Superior parietal gyrus | 31633 | -11 | -75 | 55 | 3.67 |
| Right Paracentral lobule | 29424 | -8 | -24 | 46 | 3.61 |
| Right Temporal pole: superior temporal gyrus | 13109 | -44 | 10 | -17 | 3.58 |
| Right Cuneus | 31915 | -17 | -67 | 20 | 3.51 |
| Right Postcentral gyrus | 22201 | -48 | -23 | 19 | 3.51 |
| Right Superior frontal gyrus, medial | 4440 | -4 | 52 | 25 | 3.49 |
| Right Fusiform gyrus | 36625 | -42 | -22 | -21 | 3.48 |
| Right Calcarine fissure and surrounding cortex | 39428 | -16 | -89 | -19 | 3.45 |
| Right Middle frontal gyrus | 20452 | -46 | 23 | 29 | 3.4 |
| Right Inferior frontal gyrus, orbital part | 19161 | -52 | 37 | -5 | 3.37 |
| Right Superior occipital gyrus | 31667 | -17 | -80 | 40 | 3.23 |
| Right Superior frontal gyrus, dorsolateral | 18009 | -25 | 52 | 34 | 3.13 |
| Right Superior frontal gyrus, medial orbital | 15996 | -1 | 18 | -17 | 3.11 |
| Right Inferior occipital gyrus | 35407 | -48 | -63 | -10 | 3.03 |
| Right Inferior parietal, but supramarginal and angular gyri | 23511 | -45 | -45 | 46 | 3.02 |
| Right Superior frontal gyrus, orbital part | 13938 | -25 | 31 | -18 | 2.86 |
| Right Parahippocampal gyrus | 14247 | -24 | 2 | -26 | 2.83 |
| Right Angular gyrus | 24035 | -27 | -66 | 34 | 2.82 |
| Right Precentral gyrus | 21025 | -39 | -18 | 43 | 2.75 |
| Right Middle frontal gyrus orbital part | 18981 | -33 | 58 | 3 | 2.63 |

**Supplementary Table 14.**
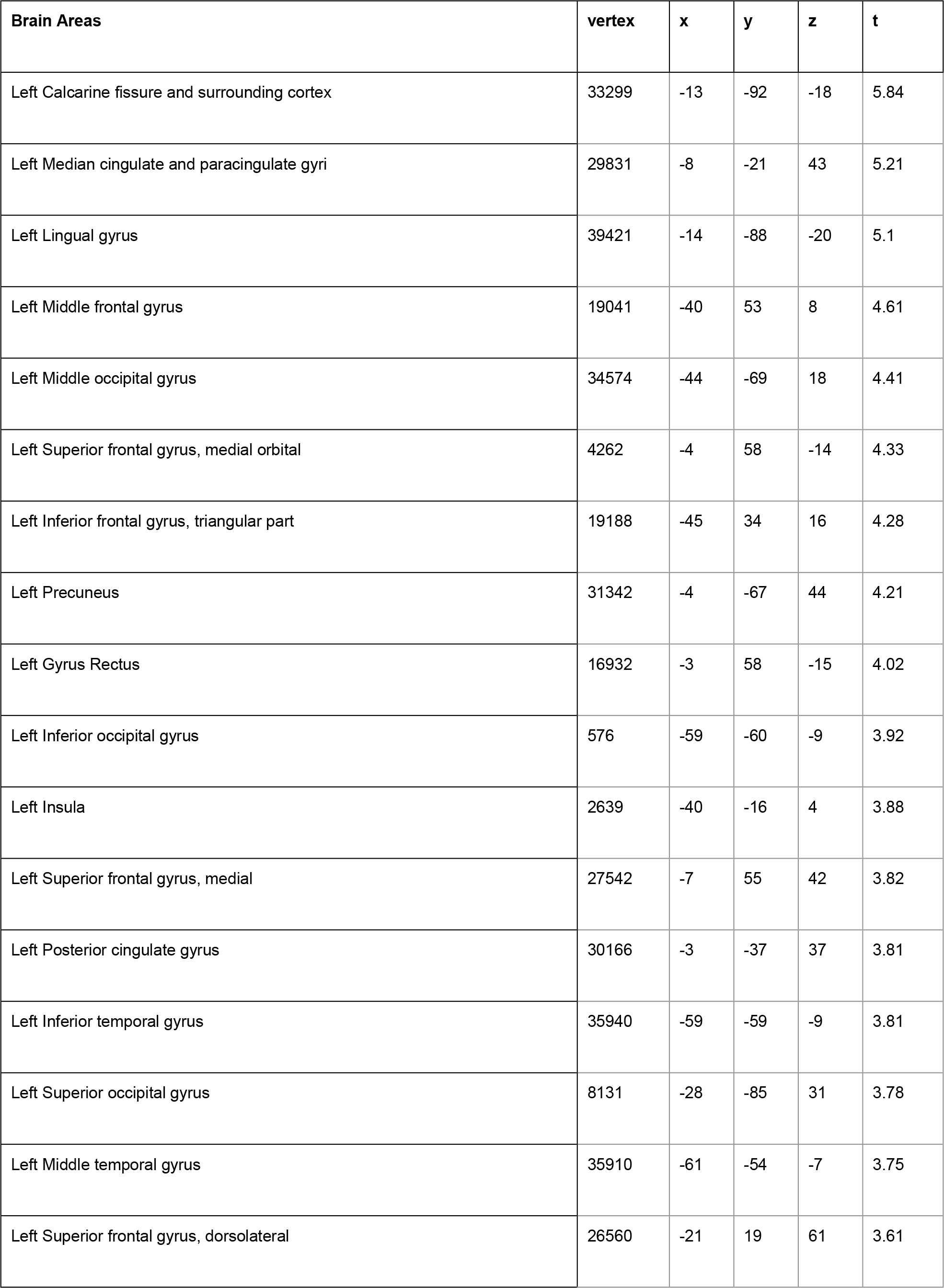
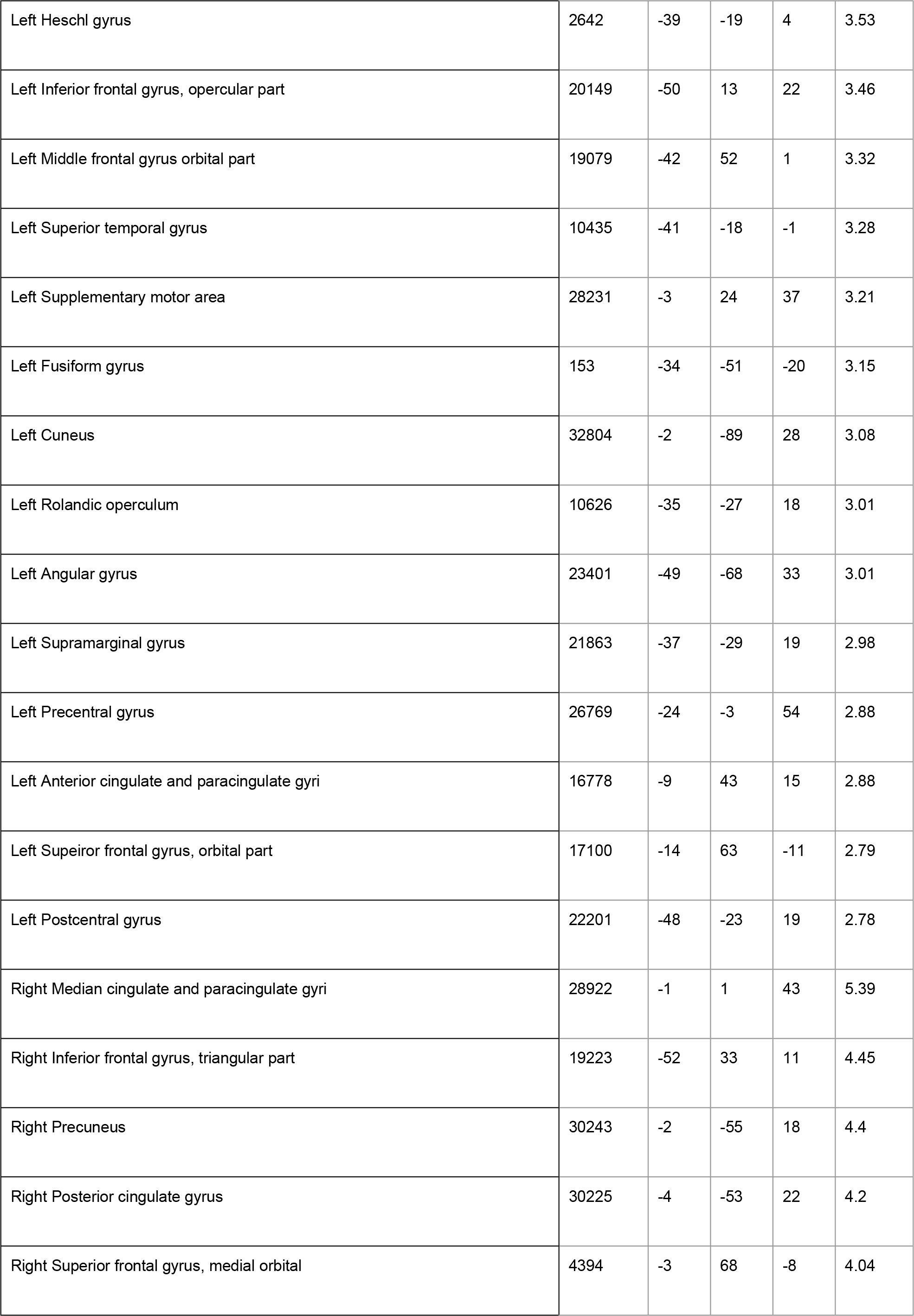
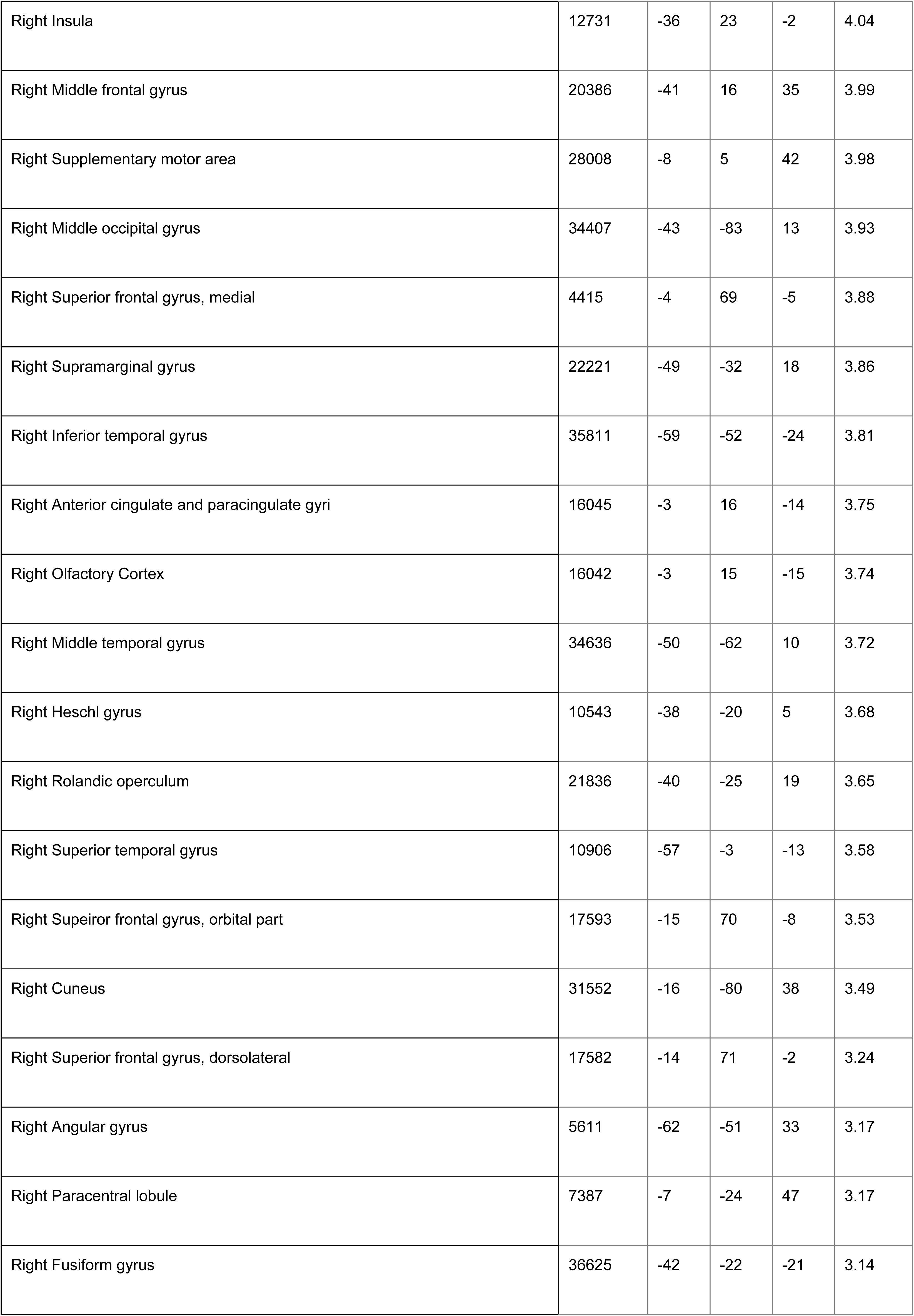
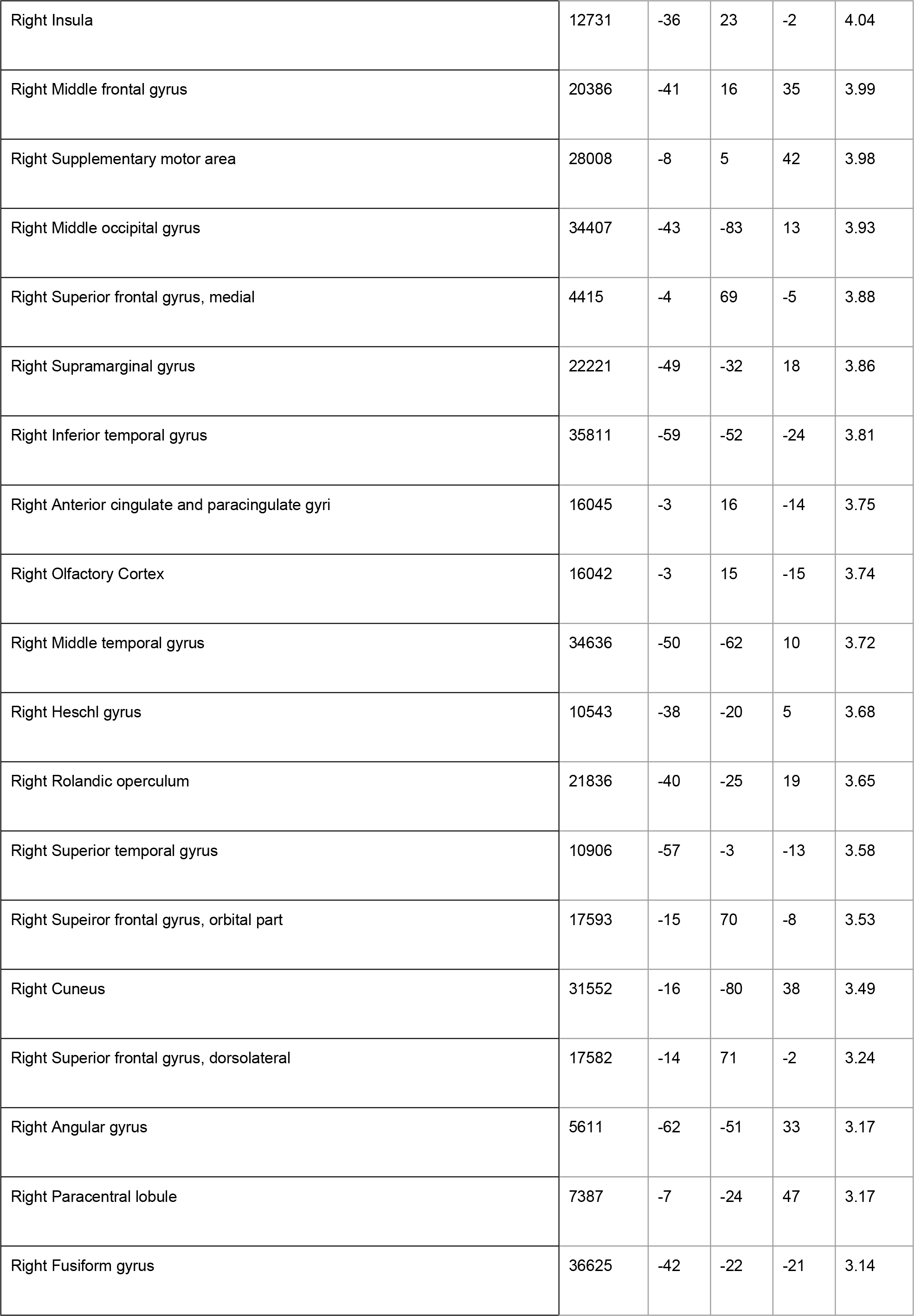
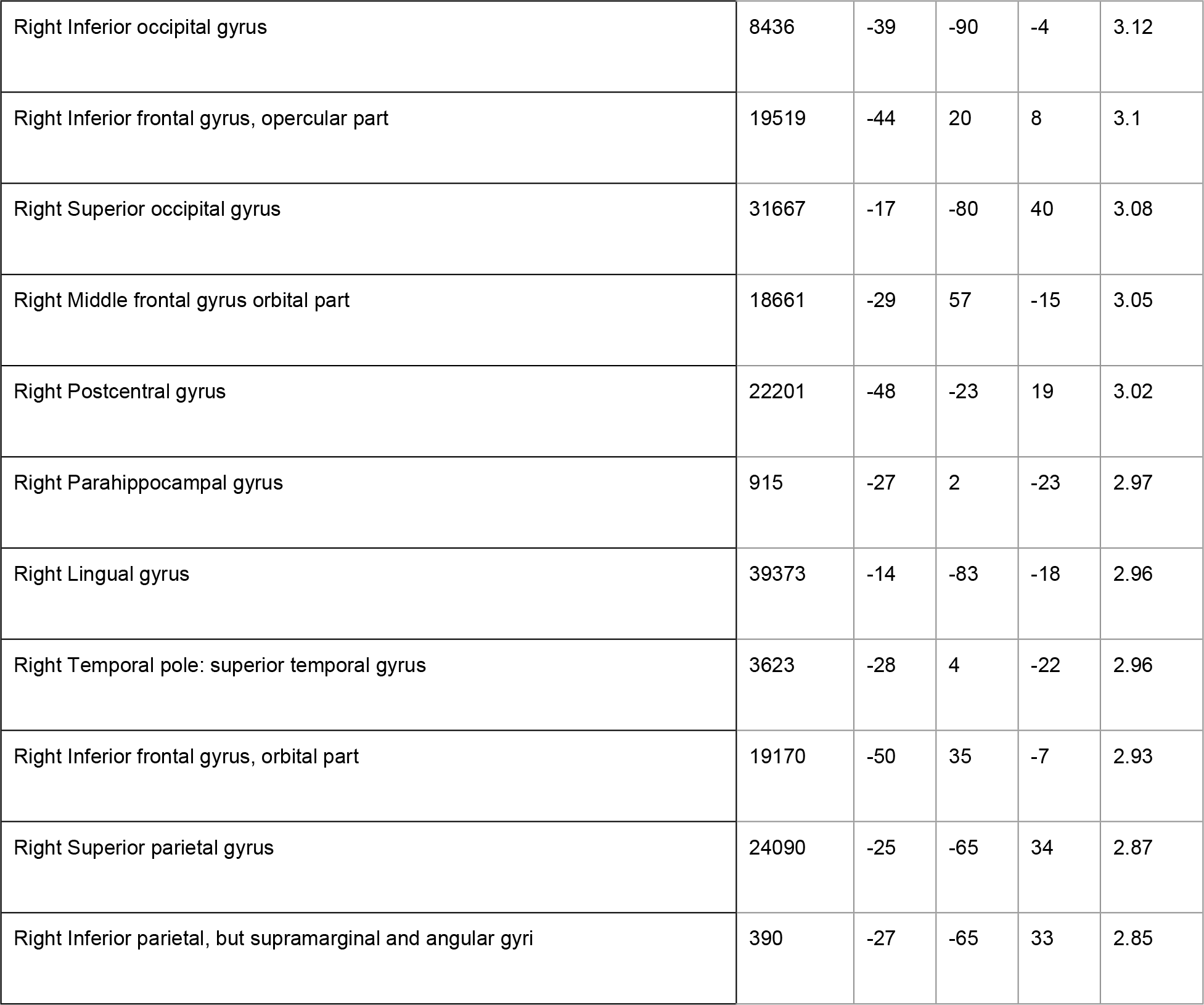
Right precommisural putamen volume significant peaks of correlation with cortical thickness for HC.

| Brain Areas | vertex | x | y | z | t |
| --- | --- | --- | --- | --- | --- |
| Left Calcarine fissure and surrounding cortex | 33299 | -13 | -92 | -18 | 5.84 |
| Left Median cingulate and paracingulate gyri | 29831 | -8 | -21 | 43 | 5.21 |
| Left Lingual gyrus | 39421 | -14 | -88 | -20 | 5.1 |
| Left Middle frontal gyrus | 19041 | -40 | 53 | 8 | 4.61 |
| Left Middle occipital gyrus | 34574 | -44 | -69 | 18 | 4.41 |
| Left Superior frontal gyrus, medial orbital | 4262 | -4 | 58 | -14 | 4.33 |
| Left Inferior frontal gyrus, triangular part | 19188 | -45 | 34 | 16 | 4.28 |
| Left Precuneus | 31342 | -4 | -67 | 44 | 4.21 |
| Left Gyrus Rectus | 16932 | -3 | 58 | -15 | 4.02 |
| Left Inferior occipital gyrus | 576 | -59 | -60 | -9 | 3.92 |
| Left Insula | 2639 | -40 | -16 | 4 | 3.88 |
| Left Superior frontal gyrus, medial | 27542 | -7 | 55 | 42 | 3.82 |
| Left Posterior cingulate gyrus | 30166 | -3 | -37 | 37 | 3.81 |
| Left Inferior temporal gyrus | 35940 | -59 | -59 | -9 | 3.81 |
| Left Superior occipital gyrus | 8131 | -28 | -85 | 31 | 3.78 |
| Left Middle temporal gyrus | 35910 | -61 | -54 | -7 | 3.75 |
| Left Superior frontal gyrus, dorsolateral | 26560 | -21 | 19 | 61 | 3.61 |
| Left Heschl gyrus | 2642 | -39 | -19 | 4 | 3.53 |
| Left Inferior frontal gyrus, opercular part | 20149 | -50 | 13 | 22 | 3.46 |
| Left Middle frontal gyrus orbital part | 19079 | -42 | 52 | 1 | 3.32 |
| Left Superior temporal gyrus | 10435 | -41 | -18 | -1 | 3.28 |
| Left Supplementary motor area | 28231 | -3 | 24 | 37 | 3.21 |
| Left Fusiform gyrus | 153 | -34 | -51 | -20 | 3.15 |
| Left Cuneus | 32804 | -2 | -89 | 28 | 3.08 |
| Left Rolandic operculum | 10626 | -35 | -27 | 18 | 3.01 |
| Left Angular gyrus | 23401 | -49 | -68 | 33 | 3.01 |
| Left Supramarginal gyrus | 21863 | -37 | -29 | 19 | 2.98 |
| Left Precentral gyrus | 26769 | -24 | -3 | 54 | 2.88 |
| Left Anterior cingulate and paracingulate gyri | 16778 | -9 | 43 | 15 | 2.88 |
| Left Superior frontal gyrus, orbital part | 17100 | -14 | 63 | -11 | 2.79 |
| Left Postcentral gyrus | 22201 | -48 | -23 | 19 | 2.78 |
| Right Median cingulate and paracingulate gyri | 28922 | -1 | 1 | 43 | 5.39 |
| Right Inferior frontal gyrus, triangular part | 19223 | -52 | 33 | 11 | 4.45 |
| Right Precuneus | 30243 | -2 | -55 | 18 | 4.4 |
| Right Posterior cingulate gyrus | 30225 | -4 | -53 | 22 | 4.2 |
| Right Superior frontal gyrus, medial orbital | 4394 | -3 | 68 | -8 | 4.04 |
| Right Insula | 12731 | -36 | 23 | -2 | 4.04 |
| Right Middle frontal gyrus | 20386 | -41 | 16 | 35 | 3.99 |
| Right Supplementary motor area | 28008 | -8 | 5 | 42 | 3.98 |
| Right Middle occipital gyrus | 34407 | -43 | -83 | 13 | 3.93 |
| Right Superior frontal gyrus, medial | 4415 | -4 | 69 | -5 | 3.88 |
| Right Supramarginal gyrus | 22221 | -49 | -32 | 18 | 3.86 |
| Right Inferior temporal gyrus | 35811 | -59 | -52 | -24 | 3.81 |
| Right Anterior cingulate and paracingulate gyri | 16045 | -3 | 16 | -14 | 3.75 |
| Right Olfactory Cortex | 16042 | -3 | 15 | -15 | 3.74 |
| Right Middle temporal gyrus | 34636 | -50 | -62 | 10 | 3.72 |
| Right Heschl gyrus | 10543 | -38 | -20 | 5 | 3.68 |
| Right Rolandic operculum | 21836 | -40 | -25 | 19 | 3.65 |
| Right Superior temporal gyrus | 10906 | -57 | -3 | -13 | 3.58 |
| Right Superior frontal gyrus, orbital part | 17593 | -15 | 70 | -8 | 3.53 |
| Right Cuneus | 31552 | -16 | -80 | 38 | 3.49 |
| Right Superior frontal gyrus, dorsolateral | 17582 | -14 | 71 | -2 | 3.24 |
| Right Angular gyrus | 5611 | -62 | -51 | 33 | 3.17 |
| Right Paracentral lobule | 7387 | -7 | -24 | 47 | 3.17 |
| Right Fusiform gyrus | 36625 | -42 | -22 | -21 | 3.14 |
| Right Inferior occipital gyrus | 8436 | -39 | -90 | -4 | 3.12 |
| Right Inferior frontal gyrus, opercular part | 19519 | -44 | 20 | 8 | 3.1 |
| Right Superior occipital gyrus | 31667 | -17 | -80 | 40 | 3.08 |
| Right Middle frontal gyrus orbital part | 18661 | -29 | 57 | -15 | 3.05 |
| Right Postcentral gyrus | 22201 | -48 | -23 | 19 | 3.02 |
| Right Parahippocampal gyrus | 915 | -27 | 2 | -23 | 2.97 |
| Right Lingual gyrus | 39373 | -14 | -83 | -18 | 2.96 |
| Right Temporal pole: superior temporal gyrus | 3623 | -28 | 4 | -22 | 2.96 |
| Right Inferior frontal gyrus, orbital part | 19170 | -50 | 35 | -7 | 2.93 |
| Right Superior parietal gyrus | 24090 | -25 | -65 | 34 | 2.87 |
| Right Inferior parietal, but supramarginal and angular gyri | 390 | -27 | -65 | 33 | 2.85 |

**Supplementary Table 15.** Right precommisural putamen volume significant peaks of correlation with cortical thickness for AD.

| Brain Areas | vertex | x | y | z | t |
| --- | --- | --- | --- | --- | --- |
| Left Superior temporal gyrus | 13033 | -44 | 3 | -15 | 4.86 |
| Left Temporal pole: superior temporal gyrus | 3278 | -44 | 6 | -15 | 4.53 |
| Left Temporal pole: middle temporal gyrus | 13285 | -48 | 13 | -30 | 4.4 |
| Left Middle temporal gyrus | 13282 | -49 | 13 | -29 | 4.36 |
| Left Inferior occipital gyrus | 33470 | -36 | -88 | -17 | 4.18 |
| Left Precentral gyrus | 20887 | -53 | -5 | 30 | 3.94 |
| Left Postcentral gyrus | 5245 | -54 | -5 | 28 | 3.76 |

